# Learning Human T Cell Behaviors through Generative AI Embeddings of T Cell Receptors

**DOI:** 10.1101/2025.06.16.660016

**Authors:** Daniel G. Chen, Yapeng Su, James R. Heath

## Abstract

T cells interact with the world through T cell receptors (TCRs). The extent to which TCRs determine T cell behavior has not been comprehensively characterized. Our Tarpon model leverages advances in generative artificial intelligence to synthesize large-scale (>1M sequences) TCR atlases across human development and diseases into actionable insights. Tarpon creates: 1) bespoke sampling functions generating realistic Ag-specific TCRs, 2) embeddings revealing CD4^+^ and CD8^+^ single-positive TCR repertoires as distinct with divergent physiochemical properties, and 3) cross-dataset mappings of T cell states that validate fetal CD4^+^ versus CD8^+^ TCR differences in adults and find fetal type I innate T cells to map to MAIT and KIR^+^ adult CD8^+^ T cells which we verify via whole transcriptome analysis. Tarpon is a resource as a reference of TCRs across human physiological states and as a computational framework to create interpretable TCR embeddings, via physicochemical associations, that have broad implications for the field.

## Introduction

T cells spearhead the cell-mediated arm of human adaptive immune response^1,2^. They execute diverse functions from immunomodulation through paracrine signaling^3–5^ to direct cell killing through their release of cytotoxic granules^6^. These actions are often guided by the T cell receptor (TCR) sequence that determines the antigen specificity of the T cell^7–10^. For example, TCRs specific for tumor-associated antigens facilitate the clearance of cancer cells^11–13^ while self-targeting TCRs can drive autoimmune disease^14,15^. Observations of T cells with similar phenotypes possessing similar TCRs in a variety of contexts raises the possibility of deterministic relationships between the TCR and a given T cell’s behavior^8,10,16–18^. Recent technological advances have fueled the creation of large TCR atlases across a diversity of human physiological contexts^19–37^. These atlases raise a unique opportunity to query if human T cell states are generally mediated by the TCR. Here we use this breadth of data to train Tarpon to be representative of human TCRs across developmental time and disease states.

Artificial intelligence (AI) can provide an effective route towards the extraction of deep biological insights from large datasets^38^. The inclusion of, for example, physicochemical properties^39,40^ or gene regulatory networks^41,42^, into AI models can imbue them with biological understandings that enhance their capabilities and yield interpretable outputs. Thus, a large-scale AI model of TCRs may help resolve whether TCR sequences influence T cell behavior *in vivo*. By explicitly designing Tarpon’s AI model with physicochemical TCR attributes (e.g. hydrophobicity, charge, etc.) we allow for its learned representations of TCRs to be human-interpretable and thus mineable by future works, such as for rational TCR design in cell therapy contexts^43,44^.

Generative AI (GenAI) models are capable of producing original data characteristic of but not necessarily from a given dataset, by learning the dataset’s underlying probability distributions^45^. GenAI has driven many recent advances in biology^46–51^, such as “inpainting” in protein design where a model paints in missing portions of a protein structure by learning the probability of structure X given known neighboring structures Y_1_…Y_N_^52^. In a similar vein, a GenAI model of TCRs may be able to *de novo* generate TCRs from any biological distribution, such as for a specific antigen (Ag). We provide a proof-of-concept for this objective by creating an array of Ag-biased TCR generation modules that incline Tarpon to only produce TCRs with high probability of binding a given Ag, an ability that opens doors for Tarpon’s application for TCR discovery efforts.

We then demonstrate Tarpon’s additional applicability by harnessing its interpretable TCR embeddings to resolve the distinct TCR repertoires that divide CD4^+^ and CD8^+^ T cell populations during thymic T cell development. By envisioning Tarpon embeddings as a “molecular fingerprint” we can validate these identified differences in adult donors, across multiple physiological conditions. Last, we unveil previously hidden heterogeneity in a rare fetal T cell population, type I innate T cells, by mapping it to adult MAIT and KIR^+^ CD8^+^ T cells, a finding that we are able to validate on the transcriptomic level. Tarpon serves a valuable community resource from its large-scale reference of TCRs from human donors of diverse physiological states to its computational framework that creates interpretable TCR embeddings. These embeddings can be leveraged for TCR discovery, biological insight mining, and cross-dataset mapping efforts. Tarpon will only grow more useful with time as TCR datasets scale and profiling technologies mature.

## Results

### Collect, Curate, Build – Creating a High-Quality TCR Reference Atlas

To create a generative artificial intelligence (GenAI) model representative of the diversity of human T cell receptors (TCRs), we collected TCR sequences from atlases comprising a breadth of human physiological states from the developing thymus to adult patients with cancer^19–37^. We curated these datasets for high-quality TCRs focusing on the third complementarity-determining region (CDR3) as it is the most commonly reported TCR feature and encodes for the highly variable region of the TCR^53,54^ (see **Methods**). We made an additional effort to collect antigen (Ag) specific TCRs restricted to major-histocompatibility-complex (MHC) class I due to the relevance of these TCRs to biomedical applications ranging from diagnostics to cell therapy efforts^55–58^. In total, we compiled 342,844 CDR3αs, 753,755 CDR3βs, and 54,883 Ag-resolved CDR3αβ pairs (**Fig. 1A**, see **Data Availability**).

**Figure 1.**
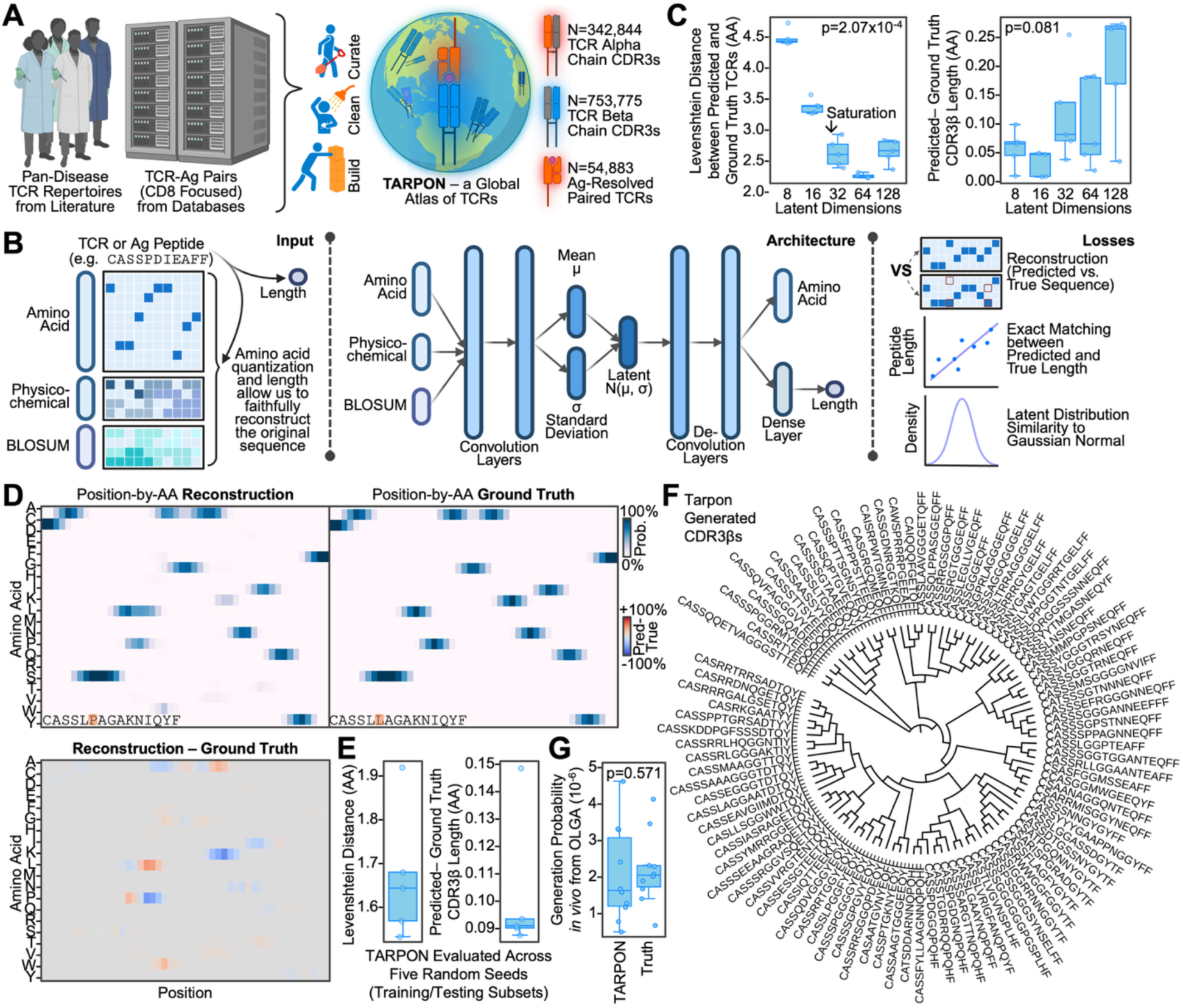
**Tarpon is a generative AI model for the embedding and accurate reconstruction of a diversity of TCRs:** A) Illustration of the curation, data cleaning, and atlas building process implemented to collect T cell receptor (TCR) sequences from a breadth of human physiological states with and without paired antigen (Ag) information. B) Illustration of Tarpon’s inputs (left) of amino acid sequence, physicochemical features, and BLOSUM62 vectors. The model architecture (center), based on a convolutional variational autoencoder Losses (right) consist of a reconstruction loss in amino acid sequence, difference in predicted versus observed sequence length, and the difference between the distribution of each latent dimension with a readily sample-able Gaussian (normal) distribution. C) Box plots illustrating how Tarpon reconstructed TCRs compare to the original sequence in terms of the metrics of Levenshtein distance (left) and sequence length (right). Both plots are in units of amino acids (AAs) and are functions of the number of latent dimensions contained within Tarpon’s model architecture (x-axis). Each dot represents one cross-validation iteration whereby a random subset of the full dataset was used for training with evaluation on the complementary test set. P-values were computed using the Kruskal-Wallis statistical test. D) Heatmap of the interpolated probability of each AA (rows) at each position in a given sequence, see color bar to the right. Columns are ordered from (left) N-terminus to (right) C-terminus. Labels on the lower left of each heatmap indicate if the TCR is the original input or reconstructed sequence. E) Box plots of how reconstructed TCRs compare to the original sequence in terms of Levenshtein distance (left) and sequence length (right), each in AA units. Each dot represents one cross-validation iteration. F) Dendrogram of 100 CDR3β AA sequences generated by Tarpon with the tree computed by the alignment algorithm MUSCLE. The tree is presented in a branch length agnostic manner and rooted based on default parameters from MUSCLE. G) Box plots of the *in vivo* generation probability, computed by OLGA, of TCRs generated by Tarpon, and TCRs in public human datasets. Each dot represents a random sample of TCRs. P-value was computed using the Mann-Whitney U test. Box plots represent median and interquartile range.

### A Generative AI Framework for T Cell Receptor Embeddings – Tarpon

Inspired by current computational approaches to TCR data, we modeled sequences as position-by-feature matrices considering three feature sets: 1) amino acid (AA) identity, 2) physiochemical attributes, and 3) BLOSUM62 vectors (**Fig. 1B left**). We accounted for variable TCR length by interpolating all sequences to a common length of 48AAs. This was the minimum length required to minimize the difference between the original and interpolated-then-extrapolated sequences. Thus, all position-by-feature matrices were interpolated to 48AAs prior to model input (**Fig. S1**). Tarpon leverages a convolutional variational autoencoder architecture to learn latent dimensions, TCR embeddings, that can regenerate the length and sequence of the originally inputted TCR (**Fig. 1B**).

Tarpon’s final architecture was composed initial and secondary convolutional layers with 256 filters, position-by-feature rule sets each, a bottleneck of 32 latent dimensions, and a 32-node dense layer to predict length (**Fig. 1B**). We confirmed these hyper-parameters as the minimum required to accurately reconstruct inputted TCR sequence and length by evaluating Tarpon variants with 2x lower to 2x higher hyper-parameter values (**Fig. S2**). As an example, we evaluated models with 8, 16, 32, 64, and 128 latent dimensions and found TCR reconstruction accuracy, measured by Levenshtein distance^59^, to saturate at 32 dimensions (**Fig. 1C**). There was no statistical difference in length prediction between model variants. Tarpon’s lighter architecture^48,60,61^ of 2M parameters affords computational efficiency to produce embeddings for 100M TCRs in just 17.45±0.55 minutes (mean±95% confidence interval) on a single GPU chip which possesses just 12.8% of the performance capacity of recently released GPUs (i.e. NVIDIA’s Blackwell GeForce RTX 5090) (**Fig. S3**).

### Tarpon Accurately Reconstructs TCRs and Generates a Diversity of Sequences

Having settled upon a robust architecture, we performed cross-validation on Tarpon using five randomized train-test splits of the entire dataset (see **Methods, Fig. S4**). Indeed, Tarpon consistently reconstructed both the sequence and length of inputted TCRs from its latent dimensions, 1.66±0.14AA difference in predicted sequence and 0.10±0.02AA difference in predicted length (**Fig. 1D-E**). For a visual example, we present how Tarpon predicts a position-by-AA matrix for a TCR not present during training that results in an extrapolated TCR CDR3β sequence nearly identical to the original, CASSL<u>P</u>AGAKNIQYF (reconstructed) versus CASSL<u>L</u>AGAKNIQYF (original) (**Fig. 1D, Fig. S5**). Similar results were observed for CDR3α (**Fig. S6**).

Thus, we present Tarpon as a scalable GenAI model for the computation of TCR embeddings that facilitate faithful TCR sequence reconstruction, and as a model that can *de novo* generate diverse TCR sequences (**Fig. 1F**) that can realistically occur in human physiological contexts, as evaluated by a well-known TCR generation probability algorithm, OLGA^62^ (**Fig. 1G**). We focused on CDR3βs in our downstream analyses as they are the most abundantly reported TCR element and thus allow for the greatest insights.

### Tarpon TCR Embeddings are Interpretable via Physiochemical Associations

Tarpon embeddings are continuous numerical representations of TCRs. We visualize these embeddings by projecting them in 2D space via Uniform Manifold Approximation and Projection (UMAP) with unsupervised clustering by the well-known Leiden algorithm (**Fig. 2A**)^63,64^. We observed mild trends with TCR length (**Fig. 2B**) and a clear stratification of TCR clusters by Tarpon latent dimensions, the 32-dimensions compose a single TCR embedding vector (**Fig. 2C**). Of note, we observed minimal correlations between the values of each latent dimension; the maximum correlation coefficient magnitude was only 0.12 (**Fig. S7A**). We highlight an example of one of the more antagonistic latent dimension pairs, 10 and 12, which had a correlation coefficient more negative than 98.19% of all other pairwise correlations (**Fig. 2C, S7B**). This suggests that Tarpon, by in large, learns a non-redundant embedding space that can still capture opposing TCR programs that may encode or associate with opposing biological programs.

**Figure 2.**
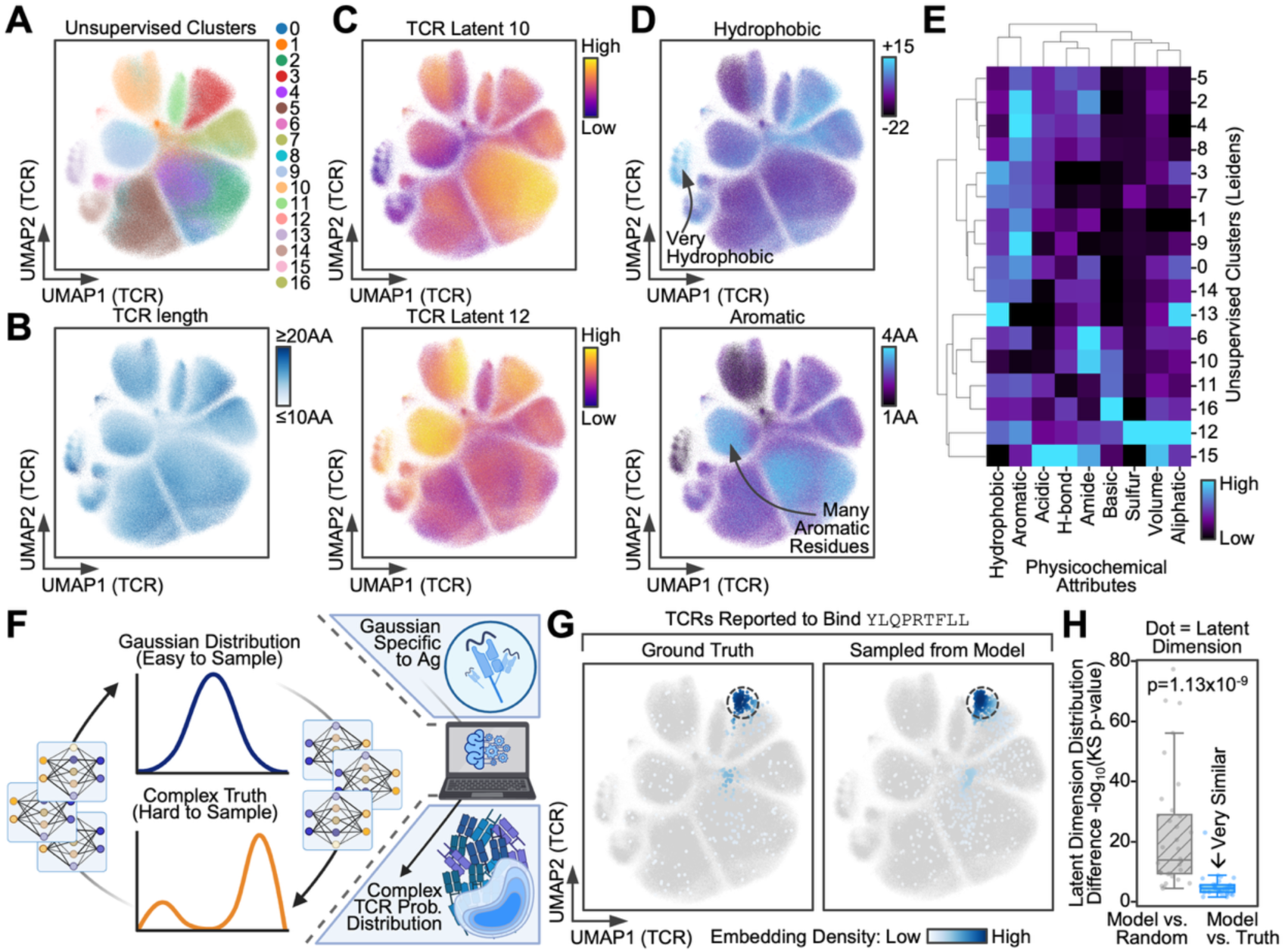
**Tarpon TCR embeddings encode for physicochemical combinatorics and distinguish antigen-specific sequences:** A-D) Scatterplot of all TCRs in our atlas with coordinates as a uniform manifold approximation projection (UMAP) of their Tarpon model latent dimensions embeddings. The colors represent (A) unsupervised clusters, via the leiden algorithm; (B) TCR length; (C) latent dimension values; and (D) physiochemical features. See legends to the right of each plot. E) Heatmap of the physiochemical attributes (columns) of each TCR cluster (rows). Color represents the average value. See legend at bottom right. F) Illustration of the AgFlow computational framework. A Normalizing Flows model was utilized to convert between easy to sample Gaussian (normal) distributions and difficult to sample complex distributions Ag-specific TCRs occupy in Tarpon latent space. Ag-specific TCRs are readily generated through this framework by sampling the Gaussian distributions, converting to coordinates in Tarpon latent space, then having Tarpon reconstruct TCR sequences from these embeddings. G) Scatterplot of YLQ-specific TCRs projected onto the UMAP embedding of all TCRs with color indicating embedding density, see legend at bottom right. YLQ-specific TCRs from experimental data (ground truth) are projected on the left UMAP, and TCRs generated from the YLQ-specific AgFlow model are at right. H) Box plot of the difference between the distribution of YLQ-specific TCRs generated from the AgFlow model compared against random TCRs (grey) and “true” YLQ-specific TCRs from experimental data (blue) in Tarpon latent space. Y-axis was quantified as -log_10_(p-value) from Kolmogorov-Smirnov tests. Each dot represents one latent dimension. P-value was computed using the Mann-Whitney U test. Box plots represent median and interquartile range.

We interrogated for the interpretability of these latent dimensions by projecting the physicochemical attributes of each sequence onto the 2D TCR UMAP (**Fig. 2D, S8**). We could indeed qualitatively observe physicochemical features forming distinct patterns on the projection. For example, there are regions of high hydrophobicity or with greater-than-average numbers of aromatic residues. We quantitatively confirmed these observations by finding TCR clusters to span a wide range of physicochemical feature combinations with clusters more similar in Tarpon embedding space to possess more similar physiochemical features, such as clusters 2, 4, and 5 (**Fig. 2E**). These tight associations between Tarpon latent dimensions and TCR physicochemical properties lends an interpretability to Tarpon embeddings that aids human understanding of the model and supports mining efforts to identify TCRs with specific attributes or combinations thereof.

### Antigen Probability Distributions are Learnable in Tarpon TCR Space

In a similar vein, the field has long sought methods for *in silico* discovery or “mining” of Ag-specific TCRs^11,65^. Achievement of this objective could significantly aid fields from cell therapy to disease diagnostics by alleviating the significant efforts it currently takes to discover and validate TCRs^58^. An *in silico* model of Ag-specific TCRs could fast-track this process through virtual screening and, via GenAI, *de novo* generation of Ag-specific TCRs. To query if Tarpon could aid in this task, we conducted a case study on TCRs specific to the SARS-CoV-2 spike antigen YLQPRTFLL (YLQ). Ag-specific TCRs often possess distinct physicochemical properties tailored to their cognate peptide^9,40,66,67^. In line with this, we confirmed YLQ-specific TCRs to have unique physiochemical attributes with increased hydrophobicity and acidity, likely to match YLQ’s own hydrophobicity and to complement YLQ’s basic residue arginine (YLQP<u>R</u>TFLL) (**Fig. S9A**). In support of this thinking, 3D TCR:pMHC structures for YLQ have confirmed interactions between YLQ’s basic residue arginine with the acidic residue aspartic acid from the TCR CDR3β sequence^68^.

Given that Tarpon’s latent dimensions innately encode for physicochemical combinatorics, (**Fig. 2D-E**), we hypothesized that Tarpon embeddings themselves would isolate YLQ-specific TCRs. Indeed, we observed certain Tarpon latent dimensions, such as 27 and 28, sharply separating out YLQ-specific TCRs from background random TCRs from our atlas (**Fig. S9B**). We used Normalizing Flows^69,70^, a flow-based GenAI model, to build upon this observation and develop a method to sample from Tarpon embeddings in a YLQ-biased manner (**Fig. 2F**). Specifically, we sought to learn a set of invertible transformations between easy-to-sample Gaussian distributions and YLQ’s complex distribution in Tarpon TCR space (see **Methods**). To test this, we train a flow-based GenAI model, which we call AgFlow, on YLQ’s ground-truth distribution in Tarpon TCR space and sample for N=1000 embeddings. When we project these embeddings back onto the UMAP we find them to occupy nearly the exact same region as ground truth YLQ-specific TCRs (**Fig. 2G**). We confirmed this qualitative observation by performing Kolmogorov-Smirnov tests between the distribution of AgFlow sampled embeddings and that of YLQ-specific and random TCRs and indeed found the model to recapitulate the true YLQ distribution (**Fig. 2H, S10**).

To further validate AgFlow, we first translated sampled embeddings into TCR sequences via Tarpon’s embedding-to-sequence engine. We verified that AgFlow model generated TCR repertoire had significant overlap with the reported YLQ-specific repertoire (see **Methods**). This suggests that the model’s recapitulation of ground truth was unlikely to have occurred by random chance. We then queried for the exact YLQ-specific sequences generated by AgFlow, and found CASSPDIEAFF to be markedly overrepresented, and indeed it is a well-known public CDR3β for YLQ^71,72^ (**Fig. S11**). This recapitulation could be due to memorization or *bona fide de novo* recognition of the dominant cognate TCR for YLQ. To test for these possibilities, we removed CASSPDIEAFF from the training data. AgFlow still generated embeddings that mapped to this sequence, even as we decreased the size of the CASSPDIEAFF-absent training data (**Fig. S12A**). We also confirmed that AgFlow does not generate this TCR when trained on random data (**Fig. S12B**). Thus, AgFlow can not only recapitulate YLQ’s distribution in Tarpon TCR space but can identify the predominant cognate TCR for YLQ without ever having seen this TCR during its training phase.

Notably, not all reported TCR-pMHC pairs in public databases will lead to T cell activation, a sizeable portion of these pairs only lead to binding without functional consequences. Thus, we sought to examine whether AgFlow generated TCRs tended to be those that led to cell activation. To this end, we leveraged a recent study that experimentally tested the T cell activation capacity of hundreds of YLQ-specific TCRs^73^. Using this validated database, we first verified that certain sequence characteristics, encoded by Tarpon latent dimensions, distinguished T cell activating versus binding YLQ-specific TCRs (**Fig. S12C**). Next, we found the TCRs most frequently generated by the AgFlow model to be those with the greatest experimental YLQ specificity (**Fig. S12D-E**). Thus, not only can Tarpon latent dimensions distinguish validated T cell activating TCRs, but our AgFlow model inherently generates TCRs with greater experimentally determined YLQ specificity at increased frequencies (e.g. 10^4^ times for a validated sequence versus once for a non-validated, per million sequences generated).

AgFlow’s success with YLQ spurred our interest in other immunogenic antigens and for those frequently reported in TCR-pMHC databases^55–57^, such as cytomegalovirus (CMV) Ag NLVPMVATV. Following the footsteps of our YLQ case study, we first interrogated for the distribution of TCRs cognate to these Ags in physicochemical and Tarpon TCR space and observed broad similarities between Ags (**Fig. S13A-D, Table S1**). Normalization against the baseline profile of TCRs in our curated atlas revealed an overrepresentation of hydrophobic TCRs and Tarpon latent dimension 27. Communal upregulation of these features may suggest the existence of an immunogenicity signature shared between TCRs cognate for immunodominant Ags (**Fig. S13E**). We then trained separate AgFlow models for each of Ag and, by in large, found the models to learn distributions more similar to ground truth than random; exceptions, such as influenza A virus (IAV) Ag GILGFVFTL, possessed ground truth distributions difficult to distinguish statistically from random (**Fig. S14A-B**). We confirmed these models to recapitulate reported TCR repertoires by finding their generated repertoires to have statistically greater overlap with ground truth than expected and having no such overlap with random repertoires (**Fig. S14C**).

Several TCR distance metrics exist^74–77^, however most necessitate information that is not always available, such as paired transcriptomes. The predominant state-of-the-art method that solely require CDR3 sequences as input is TCRdist^78,79^. We thus sought to interrogate how comparable Tarpon derived TCR distances were to those from TCRdist.

For this, we computed the distance between TCRs specific to a given Ag (target) to those reported to bind other Ags (background) where good signal presents as increased distance between target and background TCRs (**Fig. S15 left**). Tarpon recapitulated the same or better signal than TCRdist across well-reported Ags in TCR-pMHC databases (**Fig. S15 right, Table S2**). Therefore, Tarpon embeddings appear to be at least comparable to the leading CDR3-only TCR distance metric.

Thus, we demonstrate the ability for Tarpon embeddings to synergize with cutting-edge GenAI tools, through our creation of AgFlow, which leverages Normalizing Flows to learn invertible transformations between easy-to-sample Gaussians and the complex distributions Ag-specific TCRs occupy in Tarpon latent dimensions. We validate AgFlow across a wide breadth of Ags to confirm the model’s ability to learn and then generate TCRs from probability distributions in Tarpon TCR space that mimic ground truth. Through our case study with YLQPRTFLL, we further reveal AgFlow to *de novo* generate a dominant public TCR without ever having seen the sequence.

Tarpon TCR Representations Stratify CD4^+^ and CD8^+^ T Cell Fate Decision

We next asked to what extent can Tarpon TCR embeddings delineate biological phenomenon? In particular, we tested whether these embeddings could provide insights into the fundamental thymic stratification of single-positive (SP) T cells into CD4^+^ and CD8^+^ populations? Previous works have pointed to the potential role of the TCR and its downstream signaling in CD4^+^ versus CD8^+^ SP fate decision. However, a comprehensive examination of this across human development and disease states has been lacking^75,80,81^. To address this, we leveraged a single-cell (sc) TCR and RNA -seq atlas of the developing human immune system to investigate how TCR repertoires change as a function of T cell development in the fetal thymus (**Fig. 3A**)^20^. When mapped to Tarpon TCR space, we found SP T cells to not only occupy a restricted subset of the space exclusive of double-positive (DP) T cells, and also that the CD4^+^ and CD8^+^ TCR repertoires present with divergent densities (**Fig. 3B**).

**Figure 3.**
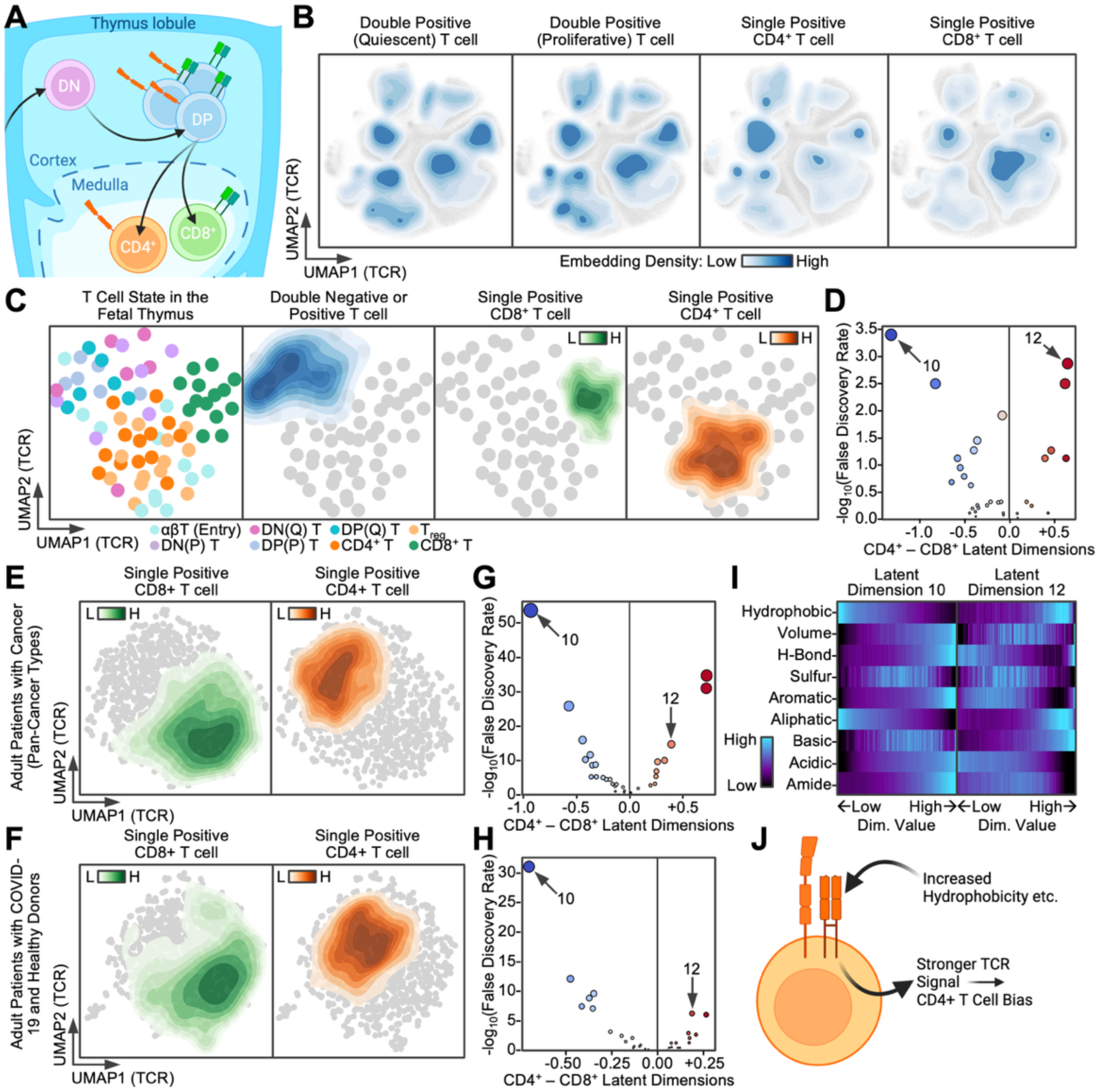
**Tarpon deciphers T cell development through a TCR lens:** A) Simplified illustration of T cell development in the fetal thymus whereby T cell progenitors differentiate into double-negative (DN) T cells, which subsequently expand into double-positive (DP) T cells. Post-selection, the DP T cells primarily become mature single-positive CD4^+^ and CD8^+^ T cells. B) Scatterplot of TCRs on the Tarpon embedding UMAP with density contours of T cells at different stages of development. See legend at the bottom. C,E,G) Scatterplots of TCR repertoires on a UMAP of their Tarpon embeddings from (C) fetal T cell development, (E) adults with COVID-19 and healthy donors, and (G) adults with cancer. Each dot represents one T cell state from one donor. Color represents T cell state on the left and density for all other plots. See the legend on the bottom and to the upper right or left of each plot. D,F,H) Scatterplots of the difference in latent dimension values between CD4^+^ and CD8^+^ populations (x-axis) against the significance of the difference (y-axis) quantified by the -log_10_(false discovery rate, FDR). Data from (D) fetal T cell development, (F) adults with COVID-19 and healthy donors, and (H) adults with cancer. I) Heatmap of the physiochemical attributes (rows) of Tarpon latent dimensions 10 and 12, for TCRs with certain Tarpon latent dimension values (columns) from low on the left to high on the right. Color represents the strength of the physicochemical feature see bottom left legend. J) Illustration illustrating a potential mechanism whereby stronger TCR signals mediated by specific sequence characteristics lead to CD4^+^ T cell specific programs and differentiation.

To test the robustness of these observations, we aggregated Tarpon embeddings by donor and T cell state and were able to reconstruct a trajectory resembling *in vivo* DP to SP T cell development, and to recover a clear bifurcation of CD4^+^ versus CD8^+^ TCR repertoires (**Fig. 3C**). We next asked whether specific latent dimensions drove the CD4^+^ versus CD8^+^ division. We identified significant enrichments of latent dimensions 10 and 12 in the CD8^+^ and CD4^+^ TCR repertoires, respectively. (**Fig. 3D**). This marked difference in TCR in fetal donors naturally asks if the same difference exists in CD4^+^ and CD8^+^ T cell populations in adults. To investigate this, we repeated these analyses in an adult cohort of patients with acute and convalescent COVID-19 with paired healthy donors and in an adult cohort of patients with diverse cancer types^19,22,23^. Remarkably, both cohorts mirrored our findings with the fetal donors. CD4^+^ TCR profiles were markedly divergent from CD8^+^ TCR profiles in a dimension 10 and 12 driven manner (**Fig. 3E-H, S16**).

For further quantitative support, we trained a cross-validated machine learning (ML) model, using logistic regression^82^, to distinguish CD4^+^ and CD8^+^ TCR repertoires using Tarpon latent dimensions. We observed statistically strong performances for all cohorts, as quantified by area under receiver-operator-characteristic (AUROC) and precision-recall curves (AUPRC) (**Fig. S17**). Notably, these findings may not be entirely unexpected as previous works have found CD4^+^ and CD8^+^ T cells to utilize different sets of TCR genes to facilitate binding to MHC-II for the former and MHC-I for the latter. As the terminal residues of the CDR3 come from the ends of TCR V and J genes, this clean classification of CD4^+^ and CD8^+^ may solely arise from the TCR gene information encoded in the CDR3. To address this, we systematically stripped away terminal CDR3 residues and assessed model performance. Even after stripping away 4AAs, longer than the 2-3AAs the V and J genes contribute^83^, we could still observe strong model performance (**Fig. S18**). Thus, Tarpon TCR embeddings can stratify CD4^+^ and CD8^+^ repertoires even when TCR gene information is stripped from the CDR3. This reveals that the divide between CD4^+^ and CD8^+^ TCRs extends beyond gene usage and to the junction sequence itself.

Given the association of the Tarpon latent dimensions with TCR physicochemical features (**Fig. 2**), we sought to identify the physicochemical rulesets that underpin the dimensions driving CD4^+^ and CD8^+^ TCR repertoire differences. This analysis revealed a stark combination of physicochemical features dividing CD4^+^ and CD8^+^ repertoires (**Fig. 3I**). For example, TCRs with high values for the CD4^+^ enriched dimension 12 either had markedly more hydrophobic or basic character relative to average (**Fig. 3I**). Thus, through a deep statistical analysis of TCR repertoires from developing T cells in the fetal thymus we reveal CD4^+^ and CD8^+^ SP fate decision to correspond with a divergent TCR repertoire that can be explained through TCR physicochemical features. These enriched features in CD4^+^ TCRs may lead to the stronger TCR signals reported to drive a CD4^+^ SP bias via GATA3 and ThPOK (also known as ZBTB7B) (**Fig. 3J**)^80,81^.

### CD4 and CD8 TCR Fingerprinting Uncover a TCR-Phenotype Relationship

Inspired by the shared latent dimensions associated with CD4^+^ and CD8^+^ TCR differences in fetal and adult cohorts, we explicitly inquired if fetal TCR characteristics that differentiate CD4^+^ and CD8^+^ T cells were the exact same as those in adulthood. We first trained cross-validated logistic regression models to distinguish CD8^+^ and CD4^+^ T cells in fetal donors. We then used these fetal data trained models to predict CD8^+^ and CD4^+^ separation in adult donors from the COVID-19 and pan-cancer cohorts (**Fig. 4A**). We repeated this analysis in reverse, training upon adult donors and predicting on the fetal cohort (**Fig. 4B**). These analyses revealed adult-to-fetal mappings to perform markedly better (AUROC=0.98±0.01, AUPRC=0.96±0.01) than fetal-to-adult mappings (AUROC=0.84±0.01, AUPRC=0.88±0.01).

**Figure 4.**
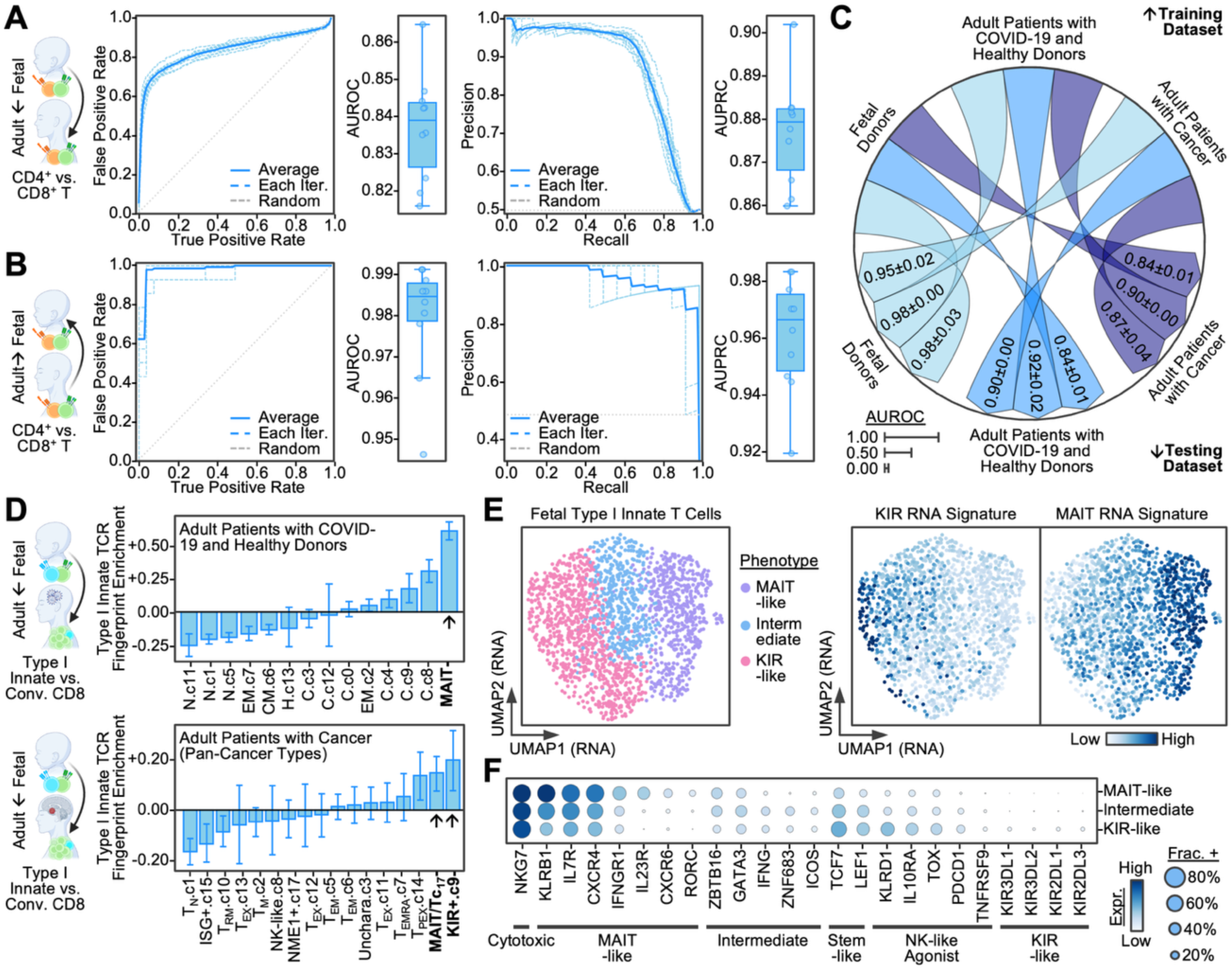
**Tarpon TCR fingerprints link T cell states across atlases:** A,B) Line plots: Receiver-operator-characteristic curves on the left and precision-recall curves on the right. Cross-validation iterations are represented by dashed blue lines, the average across iterations is in solid blue, random performance is presented as a dashed grey line. Box plots: Box plots with y-axis as the area under the receiver-operator-characteristic curve (AUROC) on the left and area under the precision-recall curve (AUPRC) on the right. Each dot represents one cross-validation iteration. Models are trained to distinguish single-positive CD4^+^ and CD8^+^ T cell classes. Upper plots are models trained on fetal data and evaluated on adult data while bottom plots are vice versa. C) Circos plot of the AUROC from cross-validated logistic regression models trained on fetal or adult cohort data, upper, and evaluated on fetal or adult cohort data, lower. Arrows point from the training dataset to the evaluation dataset with the width of the arrow representing the average AUROC value, see bottom left legend. D) Bar plots of the enrichment of fetal type I innate TCR characteristics in TCRs from T cell clusters from adult patients with COVID-19 and healthy donors (top) and adult patients with cancer (bottom). The x-axis labels are formatted to be “phenotype.subcluster”. The y-axis is the relative enrichment value, Z-score. The most strongly enriched clusters are indicated in bold font. E) Scatterplot of single type I innate T cells (each dot) on a UMAP projection of their transcriptomes. Color represents phenotype on the left (see “Phenotype” legend) and the gene expression of KIR and MAIT related genes on the right (see legend on the bottom right). F) Dot plot of the average gene expression for marker genes (columns), grouped by type I innate T cell phenotype (rows). Dot color is proportional to expression level, while dot size is proportional to the percent of cells expressing the gene. See legend at bottom right. Bar plot heights represent arithmetic mean and error bars represent 95% confidence interval. Box plots represent median and interquartile range.

To confirm this observation, we evaluated all pairwise mapping between fetal and adult cohorts and found that adult cohort trained models actually performed better on the fetal dataset than on their own. For example, a model trained on the adult pan-cancer cohort achieved an AUROC=0.87±0.04 on itself but an AUROC=0.95±0.02 on the fetal donors (**Fig. 4C**). We repeated this analysis with random forest ML models and observed a similar trend whereby adult-to-fetal mappings markedly outperformed fetal-to-adult mappings (**Fig. S19**). This suggests a significantly stronger separation of CD4^+^ and CD8^+^ TCR repertoires in the fetal donors than in adult donors even when considering healthy donors, patients with COVID-19, and patients who have cancer, with samples inclusive of those drawn from peripheral blood and diseased tissue sites.

The decreased separation of CD4^+^ and CD8^+^ TCR repertoires in adulthood, even if mild, is curious and suggests the presence of explanatory biological phenomenon. To explore this, we took advantage of our COVID-19 cohort’s deep clinical-immunophenotyping and compared patients with CD8^+^ TCR repertoires confidently assigned as CD8^+^ by the model (≥75%) to those with low confidence (<25%), i.e. incongruent with model expectations. We observed no correlation with the number of CD8^+^ T cells sequenced, excluding this technical factor as explanatory (**Fig. S20A-B**). Patients with atypical (<25% confidence) CD8^+^ TCR repertoires experienced significantly more severe acute COVID-19 and had increased abundances of inflammatory proteins, IL6 and S100A12, in their blood plasma during acute disease (**Fig. S20C-D, Table S3**). This profile suggests an impeded anti-SARS-CoV-2 response prompting us to investigate exactly how these TCRs correlate with CD8^+^ T cell phenotype. Interestingly, we found cytotoxic CD8^+^ T cells to present with the most CD8-like TCRs, and thus also the least atypical or CD4-like TCRs (**Fig. S20E**). This observation held true even when we applied CD4^+^ vs CD8^+^ TCR scoring systems derived from the cohort of fetal donors or adult patients with cancer (**Fig. S20F**).

Thus, leveraging Tarpon embeddings as a TCR lens, we unveil a fundamental divide between CD4^+^ and CD8^+^ TCR repertoires that is blurred in select adult patients who we find to present with increased disease severity. These observations prompt the usage of Tarpon embeddings to interrogate additional biological contexts and potentially unveil previously hidden connections between patient TCR repertoire and disease state.

### Cross-Dataset TCR Fingerprinting of Rare T Cell State Uncovers Hidden Heterogeneity

While our above analyses focused on conventional T cells, a notable fraction of thymus-derived T cells take on unconventional fates, such as type I innate T cells^84^. The adult equivalents for these subsets are not all well-resolved; Tarpon’s ability to genetically map T cell subsets across datasets allows us to begin to tackle this problem. Using type I innate T cells as a case study due to their intriguing cytotoxic phenotype despite being in the fetal thymus, we first confirmed this unconventional CD8^+^ subset to have TCRs distinct from conventional CD8^+^ T cells (**Fig. S21**). We then mapped its TCR fingerprint to adult T cells and observed marked enrichment in adult MAIT and KIR^+^ subsets (**Fig. 4D**). This dual enrichment prompted us to investigate if fetal type I innate T cells truly possessed such heterogeneity. Indeed, when we interrogated their whole transcriptomes, we could find this duality with one fraction of these unconventional cells appearing MAIT-like and the other fraction KIR-like based on marker genes (**Fig. 4E-G**). We confirmed the MAIT-like subset as enriched for the predominant MAIT Vα gene, TRAV1-2^85^; MAIT-associated Jα genes had less clear enrichment (**Fig. S22**). We further confirmed this subset as MAIT-like by deriving a custom gene signature for fetal MAIT-like type I innate T cells and finding it to be dominantly enriched in adult MAIT cells amongst all peripheral blood T cells (**Fig. S23**). Thus, utilizing type I innate T cells as a case study, we demonstrate how one can not only use Tarpon TCR fingerprints to map T cell states across datasets but also how we can resolve previously hidden heterogeneity, which we validate via TCR and transcriptomic means, to identify MAIT and KIR -like halves to fetal type I innate T cells.

## Discussion

Tarpon is a generative AI computational engine developed from over 1M human T cell receptors (TCRs) collected from studies across human conditions ranging from the fetal thymus to infection and pan-cancer disease in adults. Tarpon interpolates all TCR variable CDR3 domains to be of equal length and ingests those sequences largely through their physicochemical characteristics to yield a 32-dimensional embedding vector for each TCR (the latent). A single TCR can be projected onto the latent and then regenerated back into a new TCR that resembles the original within 1.7 mutations and 0.1 amino acids (AAs) in length (**Fig. 1**). We used Tarpon to address multiple fundamental questions in T cell biology, with each question leading to intriguing revelations (**Fig. 5**).

**Figure 5.**
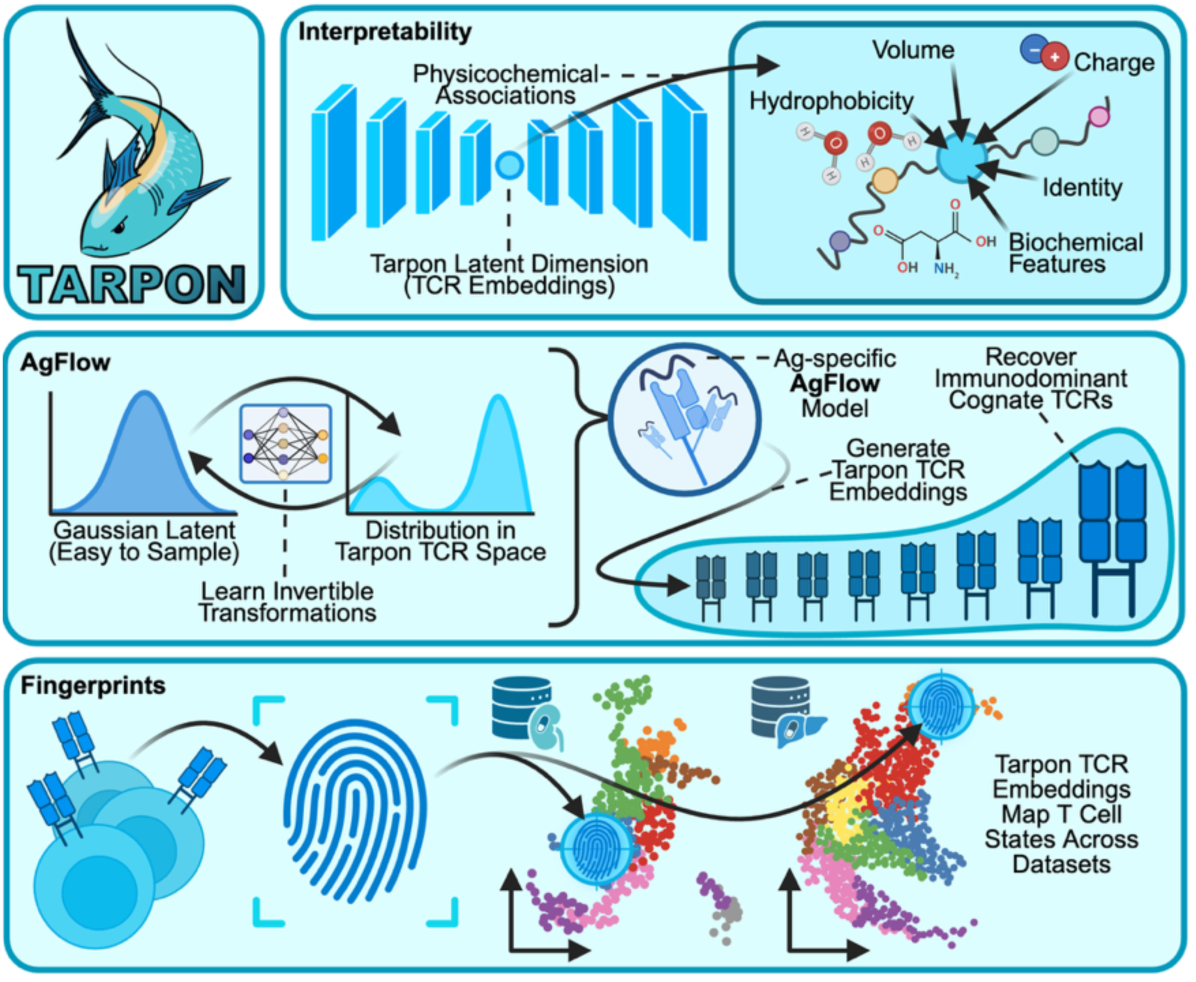
Summary of Tarpon’s Applications: Illustration depicting the architecture of Tarpon and the distinct use cases demonstrated for Tarpon in this work.

First, we queried the case of immunogenic T cell antigens (Ags), which can trigger polyclonal T cell responses, sometimes with observations of “public” TCRs that are shared across patients^86^. For several immunogenic Ags, we used Tarpon to characterize the breadth of the responding T cells and often found them to occupy a very narrow region of the full TCR atlas embedding (**Fig. S14-S15**). This supports efforts to identify Ag-specific TCRs not by exact sequence but via sequence patterns whose complexity may require models, such as Tarpon, to uncover^21,76^. For a specific immunogenic Ag from SARS-CoV-2 YLQPRTFLL (YLQ)^87^, we demonstrate how one can convert Tarpon’s distribution of TCRs to a Gaussian (normal) distribution which we use to generate a new set of Ag-specific TCRs statistically indistinguishable from experimental data (**Fig. 2F-H, S10-S12**). Remarkably, even when we removed the predominant public Ag-specific TCR from training, this same workflow could still repeatedly regenerate that “never-seen-before” TCR (**Fig. S13**). This highlights Tarpon’s capacity to generate new TCRs against target antigens, which sets it apart from most other TCR prediction algorithms, which are designed to interrogate for molecular similarities between TCRs^21,88,89^. Tarpon may ultimately provide guidance for “synthetically” engineering new TCRs against target antigens. For reported YLQ-specific TCRs, we showed that Tarpon could resolve TCRS that were capable of T cell activation relative to those reported to bind yet not experimentally identified as triggering activation^73^. Whether Tarpon can serve as a tool that can enable such discrimination more generally is currently unresolve, but will likely be increasingly achievable as multi-modal datasets measuring TCR and single-cell phenotype become increasingly available^8,10,18,90,91^.

We next queried whether Tarpon could resolve biological phenomena by applying it to characterizing T cell development within the fetus. Using just the CDR3β chains, Tarpon readily distinguished multiple steps in T cell development, including near complete separation of T cells into the single positive CD4^+^ and CD8^+^ classes (**Fig. 3**), as well as separation of type I innate CD8^+^ T cells from conventional CD8^+^ T cells. Even when major-histocompatibility-complex (MHC) specificity determining V and J gene segments^92,93^ of the CDR3β chain were removed, the remaining D gene segment and NP-nucleotides could still resolve the two T cell classes (**Fig. S19**). Notably, the same latent dimensions that were most important in resolving fetal TCRs also resolved CD8^+^ and CD4^+^ populations in adult COVID-19 and pan-cancer cohorts (**Fig. S17**). A similar projection of type I innate T cells onto adult cohort data surprisingly revealed those T cells to map to adult MAIT and KIR^+^ CD8^+^ T cells and, indeed, we subsequently found fetal type I innate T cells transcriptomes to split cleanly into MAIT-like and KIR^+^ clusters (**Fig. 4E-F, S23-S24**). Thus, at least for certain T cell states, there exists a deterministic-like relationships between genotype, TCR, and phenotype.

An advantage of developing Tarpon with datasets that span such a broad range of human biology is that one can ask similar questions, but in multiple directions – somewhat similar to a Bayesian approach – and this reveals new biology. For example, Tarpon TCR rulesets derived from the fetal cohort could be used to query one of the adult cohorts, or vice versa (**Fig. S20**). This approach suggests the fetal environment to possess well-followed rules that are partially lost in adulthood. A key contributor to, or consequence of, this loss was the emergence of “CD4-like” CD8^+^ T cells whose relative excess was seen in a subset of patients with COVID-19, and associates with severe disease, as measured through multiple metrics (**Fig. S21**). The heterogenous rise of these atypical cells in adults urges for the uncovering of factors that lead to their development, such as age, sex, and ancestry. It is likely that other health consequences exist for these cells in other disease contexts, which can be explored by future works. For type I innate T cell subtypes, we resolved some TCR gene usage differences between fetal and adult cohorts, but whether those differences translate into functional or health outcomes was not resolved for those rare cell types. With additional datasets from diverse cohorts, the resolving power of Tarpon will only increase to, hopefully, further our deciphering of T cell behavior as dictated by the TCR.

A primary limitation of this work was the size of the datasets. CDR3β chains were by far the most abundant and were, thus, employed for downstream biological investigations. CDR3α chains were only about half as abundant, and, while training Tarpon on that data yielded consistent results, the resolving power was proportionately less. The dataset of T cell antigens was an additional six-fold reduced in size which permitted very little resolution of new biology. Future technologies that more fully incorporate the TCR, such as including CDRs 1-2, will increase the amount of TCR information available for models like Tarpon to learn and to ultimately provide health-relevant predictions. In addition, the majority of our findings were based on high-resolution multi-omic datasets that simultaneously measured single-cell transcriptome and TCR information. This pairing allowed us to directly quantify links between T cell genotype and phenotype. As these multi-omic datasets become more abundant, the strength and contexts in which this genotype-phenotype link exists will become easier to resolve and characterize.

## Methods

### Curation of Diverse TCR Sequence Data

TCR, namely CDR3, and antigen (Ag) sequences were gathered from public TCR peptide-MHC (pMHC) databases (VDJdb, McPAS, and IEDB) and from atlases derived from literature^19–37,55–57^. We sought to curate TCR sequences from a diverse series of atlas. In detail we included: tumor-infiltrating lymphocytes (TILs) from patients with melanoma^27,28,34^, those found in patient with inflammatory bowel disease^30^, TILs in patients with lung cancer^24,26,37^, large granular lymphocyte leukemia T cells^33^, patients infected with HIV^32^, T cells reactive to *M. tuberculosis*^21^, pan-cancer atlases of T cells^19,29,35^, fetal donors^20^, patients with glioblastoma^31^, patients receiving immunotherapy for pancreatic cancer^25^, and patients with basal or squamous cell carcinoma^36^. Sequences from patients with acute and long COVID-19^22,23^ were not used for training due to overfitting but all cleaned sequences were utilized for downstream analyses.

### TCR Sequence Cleaning and Encoding

CDR3 sequences were cleaned as with previous works^94,95^ by adding a cysteine to sequences without leading cysteines, as this is likely due to incomplete CDR3 reconstruction. Sequences were filtered for nominal lengths, that is at least eight and with maximum length of 24 amino acids (AAs). Any sequences with non-standard amino acid lettering (e.g. “B”, “J”, “O”, “U”, “X”, “Z”) were removed. Sequences were deduplicated to avoid a particular sequence dominating model training. Sequences were encoded by amino acid identity in a one-hot-encoding manner; biological, chemical, and physical properties^94^ were encoded including hydrophobicity, volume, H-bond capabilities, and presence of sulfur, aromatic, aliphatic, basic, acidic, or amide containing side groups; and BLOSUM62^96^ was utilized to derive an additional embedding as done by certain current works^97^. BLOSUM62 values were divided by five to get the majority of embedding values between zero and one. As sequences are of variable length, we interpolated these sequences to the same length by “stretching” each peptide to 48 AAs. This length was the shortest that achieved a minimal error when interpolating a given TCR sequence to said length then extrapolating back to the original length.

### Model Architecture and Hyperparameter Determination

Our model, Tarpon, was based on a convolutional variational autoencoder with an implementation similar to previous works^98^. That is, two one-dimensional convolutional layers, a flattening layer that feeds into one dense layer to determine mean and the other to determine log-variance, the reparameterization trick samples values (z-coordinates), which are fed into a dense layer that then undergoes two convolutional transpositions which leads to the reconstruction of the interpolated position-by-AA identity matrix. An additional dense layer takes the intermediate input of the decoding to predict sequence length. All parameter values are available in detail on GitHub (see **Code Availability**).

We searched for ideal model parameters by adjusting each individually, keeping all others frozen, and examining the effect the parameter adjustment (two fold below and above the final parameter value) has on model performance as quantified by Levenshtein distance between predicted and true sequence as well as distance between predicted and true sequence length. Each search was performed in a cross-validated fashion whereby each analysis was repeated five times, each with a different random seed and thus different training and testing set, to ensure the robustness of the results. Evaluation of the finalized model architecture was also performed in a cross-validated fashion.

### Evaluating TCR Generation Probabilities

T cell generation probabilities were calculated using published algorithm OLGA^62^. Generation probabilities of the model were computed by first generating a random sample of 100 z-coordinates from Tarpon latent space then converting these coordinates into sequences via our decoder function. We then compared this with OLGA computed generation probabilities of a random set of 100 sequences from our curated set of real human TCR sequences. This was repeated 10 times to ensure result robustness.

### Learning Probability Distributions via Normalizing Flows

We leverage a public implementation of Normalizing Flows (RealNVP)^99^ which learns to map Gaussian (normal) distributions to complex distributions. In our context, this entails mapping easy-to-sample Gaussian distributions to the complex distribution TCRs cognate to a given Ag take in Tarpon latent space. Training entails 10,000 epochs on all 32 Tarpon latent dimensions of TCRs specific to a given Ag from public databases (aforementioned VDJdb, McPAS, IEDB). We visually confirm that the model, which we name AgFlow, has learned the proper distribution by randomly sampling 1,000 z-coordinates from Tarpon latent space and mapping onto our global TCR UMAP. In detail, this is done by first sampling from a Gaussian and using the AgFlow model to convert to Tarpon latent space and then using a k-nearest-neighbors regressor (with k set to five) to map z-coordinates onto the UMAP coordinates.

In our hold-out experiments, we removed a given TCR sequence from the training data and trained a new AgFlow model and tested for generation of the held-out sequence. In our efforts comparing to experimental data, we sampled one million YLQ-specific TCRs from our AgFlow model trained on TCRs cognate to the immunogenic SARS-CoV-2 spike antigen YLQPRTFLL (YLQ). We then quantified generation frequency as the number of times a given TCR was generated by the AgFlow model in those million sequences generated, i.e. how many times the model generated a z-coordinate that mapped to a given TCR sequence.

When computing for the difference between latent dimension distributions of model, generated, and random sequences we utilized the Kolmogorov-Smirnov tests which tests the significance between two distributions. For our context this entailed comparisons of the distribution of model generated Ag-specific TCRs in latent dimension one, for example, versus those occupied by ground truth Ag-specific TCRs. In testing for the significance of overlap between model generated TCR repertoires and ground truth repertoires, we utilized a hypergeometric test and utilized p<0.05 as a cutoff.

### Latent Dimension Physicochemical Profiles

To generate physicochemical profiles for a given latent dimension, we 1) rank ordered TCRs by their score in a given latent dimension, then 2) binned TCRs into percentiles by this ranking, 3) determined the average value for each physicochemical characteristic in each of these bins, which were used to 4) plot heatmaps of how physicochemical characteristics change as a function of latent dimension value.

### Machine Learning Transfer of TCR Profiles

We generated TCR profiles for each donor and each cell state. That is, one observation consists of the average Tarpon latent dimension values for TCRs in one T cell state from one sample, e.g. “effector memory T cells in COVID-19 patient #3”. We utilized these profiles for TCR embeddings and for the training and evaluation of machine learning models to predict T cell state from TCR profiles. These machine learning models included logistic regression and random forest models (with the latter having varying number of estimators). All models were cross-validated, meaning they were trained with ten different random seeds all with distinct training sets and complementary testing sets.

Models were evaluated using area under receiver-operator-characteristic curve (AUROC) and area under precision-recall curve (AUPRC) metrics on these held-out testing sets to ensure model robustness with a variety of training data. To track TCR fingerprints across datasets, we trained ML models on a given dataset, e.g. fetal donors, and then predicted on a distinct dataset, e.g. adult patients with cancer. Model outputs were normalized via Z-scores and then utilized to compare TCR similarity with cell states in the training dataset.

Prediction of CD4^+^ and CD8^+^ T Cell Type from Trimmed CDR3 Sequences

CDR3 sequences utilized for model training were trimmed by one, two, three, and four AAs form either side totaling to two, four, six, and eight total AAs trimmed from each sequence. The trimming of these sequences allow for progressive removal of V, leading, and J, ending, gene contributions to the CDR3 sequence. Tarpon models were trained as described above with these trimmed sequences and inference was performed on all trimmed sequences. Trimmed sequence latent space embeddings were then utilized for CD4^+^ versus CD8^+^ TCR repertoire classification as described above in the “Machine Learning Transfer of TCR Profiles” section.

Analysis of CD8^+^ T Cells with Atypical TCRs

Machine learning models trained to distinguish CD4^+^ from CD8^+^ T cells in patients with COVID-19 and healthy donors^22,23^ was utilized to derive per-TCR resolution scores. These scores were utilized to isolate patients with larger fractions of CD8^+^ T cells that had CD4-like TCRs, i.e. they were predicted to be more CD4 than CD8 like. The clinical characteristics of these patients were extracted from previous works to compare disease severity, i.e. World Health Organization (WHO) ordinal scale (WOS). Samples received from patients immediately when they were diagnosed with COVID-19 were utilized for patient classification and analysis. Plasma proteome samples were utilized for additional characterization of patient populations with Mann-Whitney U tests being utilized for differential protein abundance testing. We also scored single CD8^+^ T cells to identify the cell population that may be responsible for bearing a CD4-like TCR signature. We repeated this analysis with models trained on fetal donors as well as patients with cancer to ensure the robustness of our results.

### Whole Transcriptome and TCR Analysis of Fetal Type I Innate T Cells

Machine learning models were trained as per above to distinguish fetal type I innate T cells from conventional CD8^+^ T cells. Cross-validated models were utilized to predict on adult cohort data from patients with COVID-19, multiple cancers, and healthy donors^19,22^. Transcriptomic signatures enriched in the populations with high fetal type I innate T cell TCR signatures, adult MAIT and KIR-like states, were plotted on fetal type I innate T cell single-cell whole transcriptome data^20^. These signatures were computed by deriving smoothed gene expression matrices via MAGIC^100^ with the MAIT-like signature defined as *RORC, KLRB1, CXCR6,* and *IL23R*^101–104^ and the KIR-like signature was defined via the inhibitory KIR genes (*KIR2DL1, KIR2DL3, KIR3DL1,* and *KIR3DL2*)^105^.

The embedding for fetal type I innate T cells was completed by extracting labeled cells from a public pan-fetal immune atlas^20^; normalizing counts to log, natural base, counts per million plus one; utilizing the given scVI^46^ embedding with k=30 nearest neighbors to construct a k-nearest-neighbors graph; and then generating a UMAP based on said kNN. Unsupervised clustering was performed using the leiden^64^ algorithm with a resolution set to 0.9. Clusters were grouped together based on gene expression values, e.g. the signatures above, leading to clusters 0, 1, and 5 being assigned KIR-like; 2 and 6 assigned intermediate; and 3, 4, and 7 being assigned MAIT-like.

TCR gene enrichment for the clusters was computed by computing for the frequency of a given gene in either KIR or MAIT-like fractions of type I innate T cells and subtracting from that the average frequency of said gene in all fetal type I innate T cells. This analysis can be repeated to directly compare the frequency of a given TCR gene in KIR versus MAIT-like portions, i.e. subtract the frequency of said gene in the KIR-like fraction of type I innate T cells from the frequency in the MAIT-like fraction.

### Quantification and statistical analysis

Densities were calculated via contours, via seaborn^106^, or on a per-observation resolution via the stats.gaussian_kde by scipy^107^; correlations and p-values (primarily Mann-Whitney U test unless otherwise specified) were all calculated using scipy. The specific statistical test utilized to generate p-values are specified in Methods and Figure Legends. Multiple hypothesis testing correction was completed using the Benjamin-Hochberg false discovery rate (FDR) via statsmodels’s stats.multitest.fdrcorrection function^108^. kNN graph^109^, leiden^64^, UMAP^63^ calculations were completed using implentations by Scanpy^110^. All deep learning models were implemented using pytorch^111^ and all classical machine learning models were implemented using scikit-learn^109^. Phylogenetic analyses were done by computing sequence alignment and trees via MUSCLE^112^.

## Data Availability

All data for this manuscript was derived from public sources^19–37,55–57^. Processed versions of these data available at https://data.mendeley.com/preview/y3nfmh7nkb?a=f6f032f2-3acc-4c17-9822-e8514e75a343.

## Code Availability

Tarpon is available as a python package at https://github.com/danielgchen/Tarpon/. Scripts (stored in the repository as python notebooks) to reproduce the work are available at https://github.com/danielgchen/reproducibility_tarpon.

## Acknowledgements

We acknowledge the Heath lab for their support throughout this work. We thank Dr. Chenqu Suo and Prof. Sarah Teichmann at the Cambridge Stem Cell Institute, as well as Rachel X. Shi and Tiara Schwarze-Taufiq for their insightful discussions on this work. We appreciate Rachel H. Chae for iteration on graphic design. Figure cartoons were created with BioRender.com.

## Author Contributions

Conceptualization: D.G.C.; Methodology: D.G.C.; Investigation: D.G.C.; Resources: J.R.H.; Writing -Original Draft: D.G.C., J.R.H.; Writing - Review & Editing: D.G.C., Y.S., J.R.H.; Visualization: D.G.C.; Supervision: Y.S., J.R.H.; Funding acquisition: J.R.H.

## Supplementary Figures

**Figure S1.**
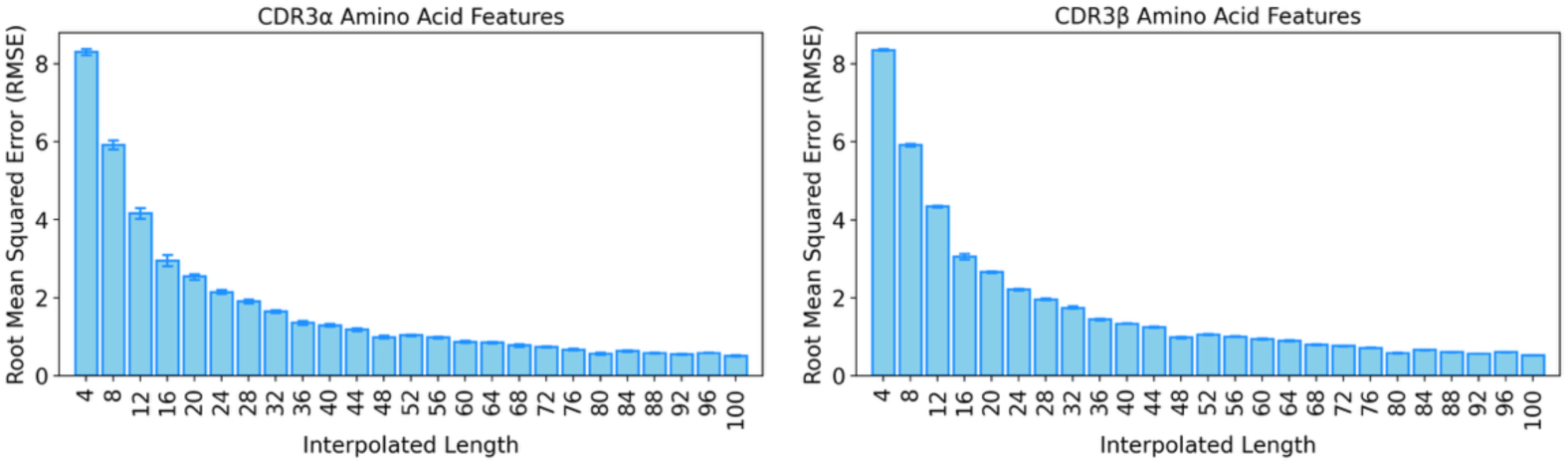
Determining an appropriate interpolation length to normalize variable TCR sequence lengths: Bar plots of root mean squared error (RMSE) between original and reconstructed, interpolated-then-extrapolated, position by amino acid (AA) feature (AA identity, physiochemical attributes, BLOSUM encoding) matrices using different lengths for interpolation. Sequences (N=100) were randomly sampled five times to generate error bars and estimates of arithmetic mean. Bar plot heights represent arithmetic mean and error bars represent 95% confidence interval.

**Figure S2.**
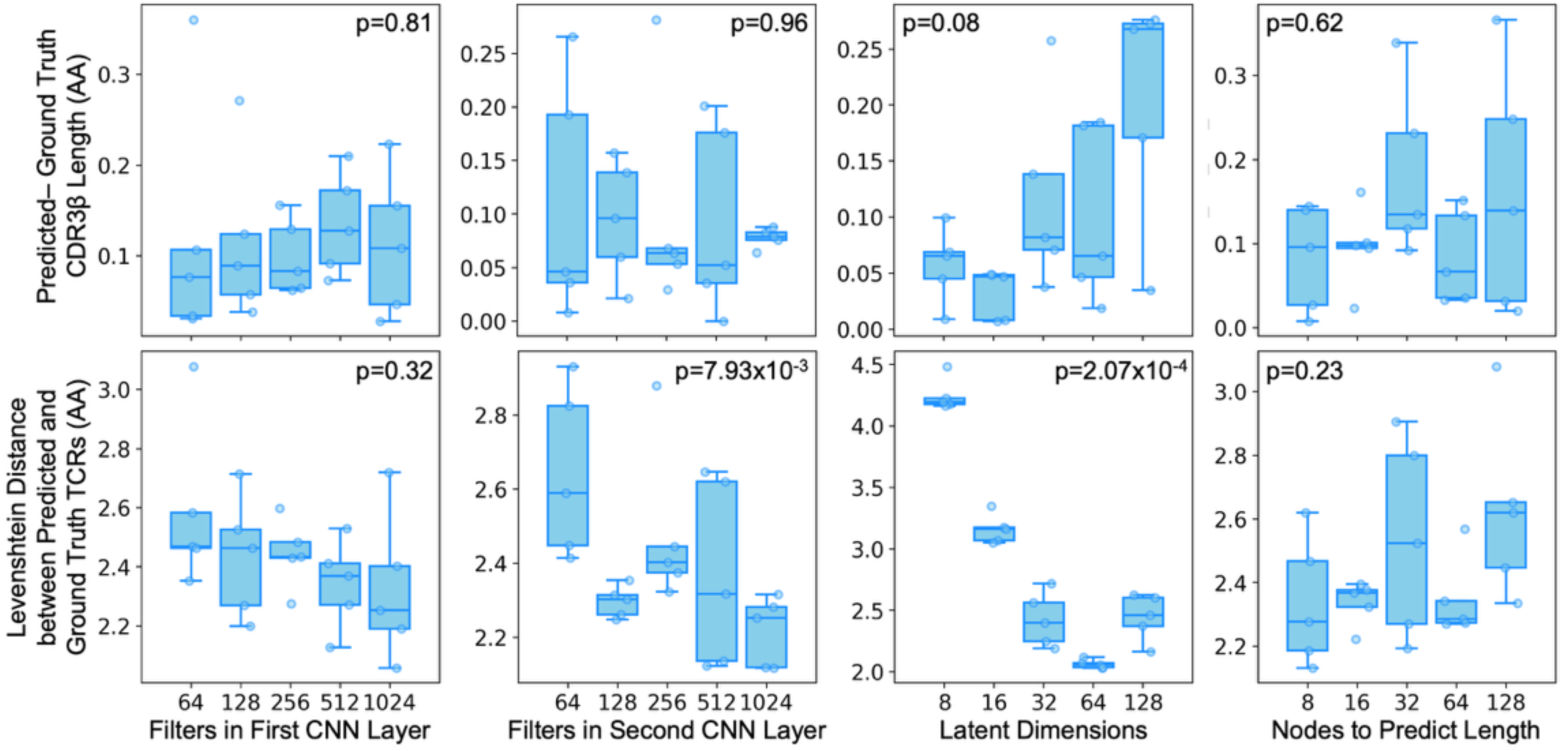
Hyperparameter grid search for Tarpon model architecture: Box plots of how reconstructed TCRs by Tarpon compare to the original sequence, on the y-axis, in terms of Levenshtein distance (left) and length (right) both with units of amino acids (AAs) as a function of various hyperparameters contained within Tarpon’s model, on the x-axis. Each dot represents one cross-validation iteration, a random subset of the dataset is used for training and evaluation of the aforementioned metrics is computed on the complementary test set. P-values were computed using the Kruskal-Wallis test. Box plots represent median and interquartile range.

**Figure S3.**
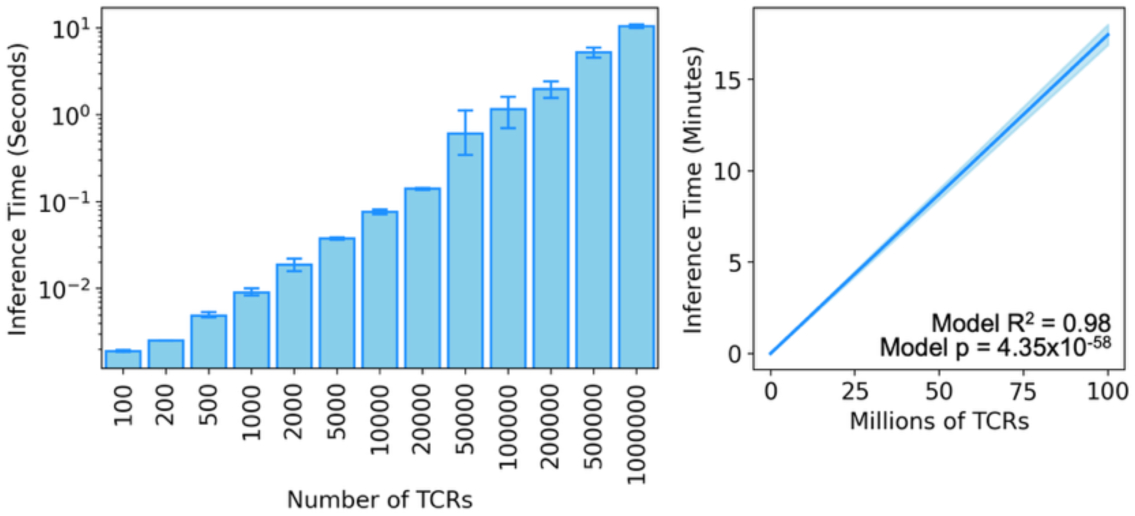
Tarpon inference time: Left: Bar plots of the inference time of Tarpon for varying numbers of TCRs. Right: Line plot of the Tarpon inference time versus numbers of TCRs (in millions). The mean is represented by a solid blue line and 95% confidence interval as a blue shade. Bar plot heights represent arithmetic mean and error bars represent 95% confidence interval.

**Figure S4.**
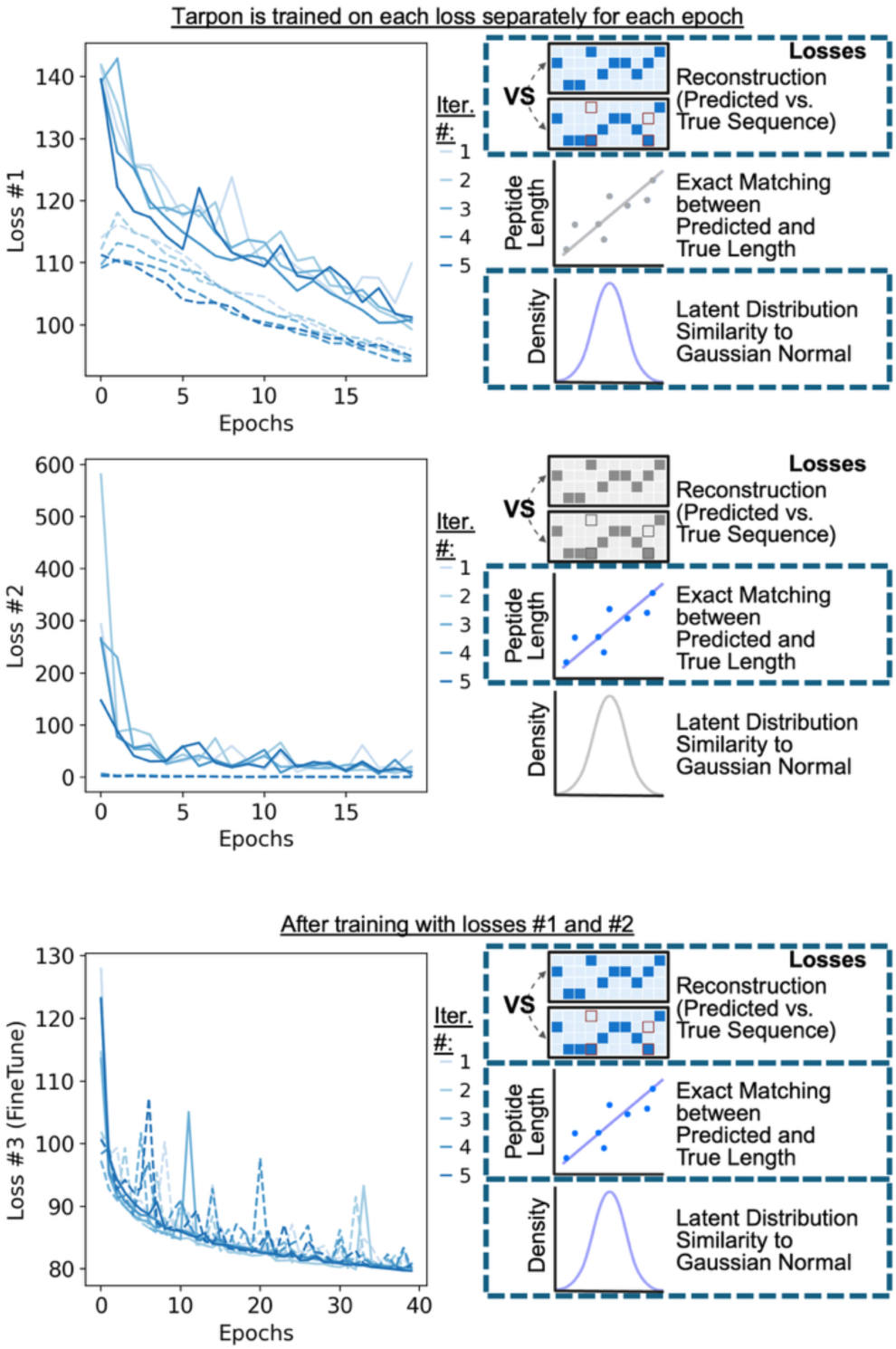
– Model losses during training: Line plots of losses during Tarpon training. Five cross-validation iterations are plotted as different colors. Training datasets are depicted as solid lines and validation datasets are dashed lines. Losses utilized in the line plots are indicted with dashed borders to the right of each plot. Losses #1 and #2, reconstruction and length, were trained in an alternating fashion for 20 epochs before all three losses were trained for an additional 40 epochs.

**Figure S5.**
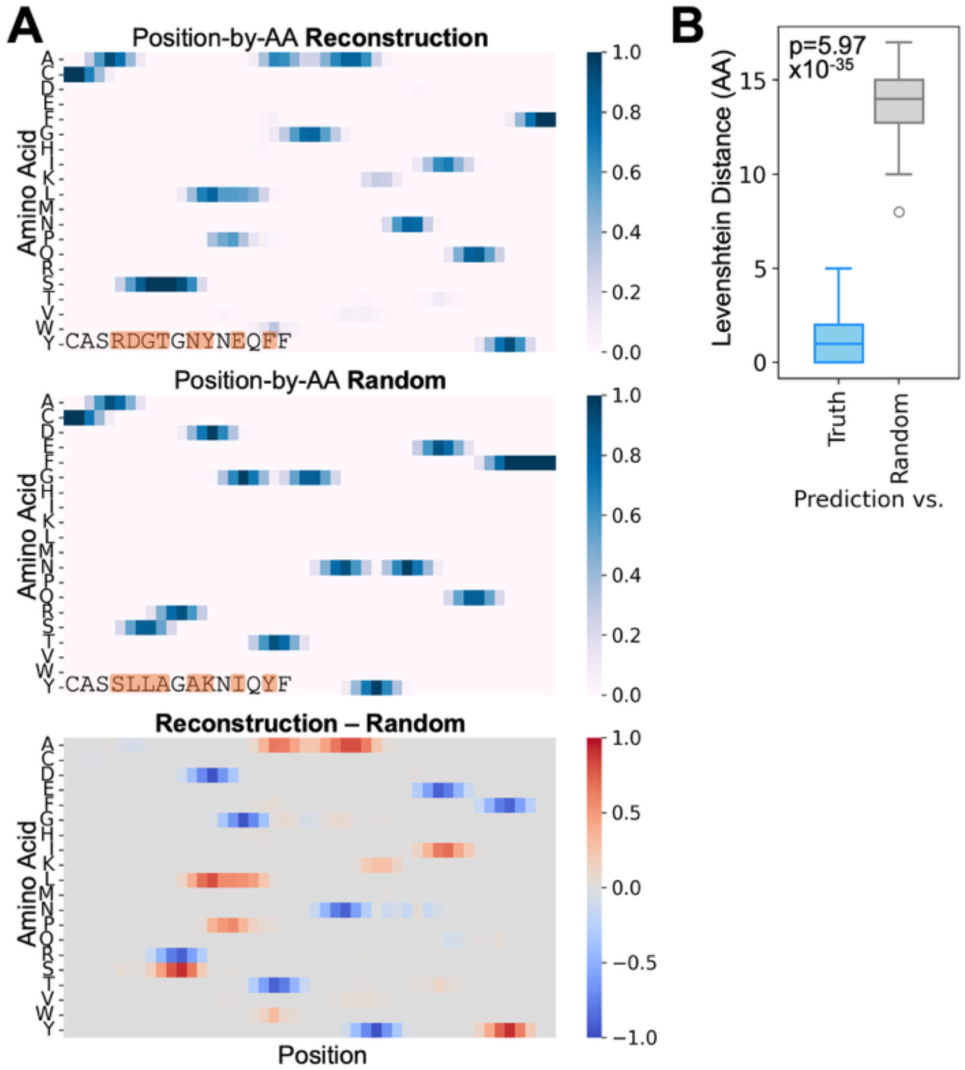
Comparison of reconstructed TCRs with random and original TCRs: A) Heatmap of the interpolated probability for each AA (rows), at each position in a given protein sequence (columns). The columns are ordered from N-terminus on the left to C-terminus. Random or reconstructed TCR sequences are labeled on the lower left of each heatmap. B) Box plots of the Levenshtein distance comparisons of original versus reconstructed TCRs (left) and random versus reconstructed (right). P-value was computed using Mann-Whitney U test. Box plots represent median and interquartile range.

**Figure S6.**
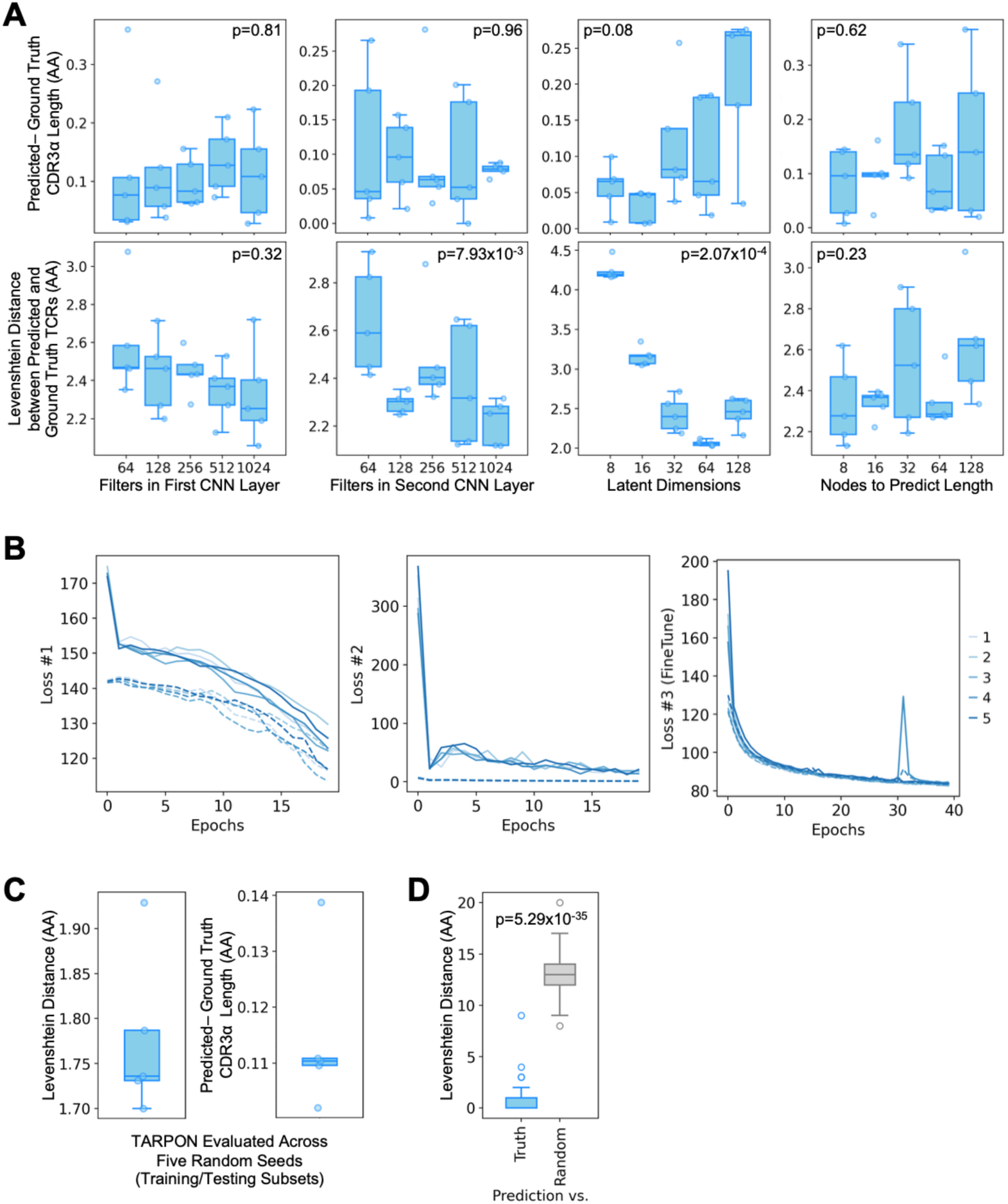
Replication of Tarpon hyperparameter grid search, training, and evaluation with CDR3α sequences: A) Box plots comparisons of how reconstructed TCRs by Tarpon relate to the original sequence with respect to length (upper) and Levenshtein (lower) distance, both with units of amino acids (AAs), and as a function of the various hyperparameters contained within the Tarpon model (x-axis). Each dot represents one cross-validation iteration. A random subset of the dataset is used for training and evaluation of the Tarpon parameters and is computed on the complementary test set. P-values were computed using the Kruskal-Wallis test. B) Line plots of losses during Tarpon training with five cross-validation iterations, different colors see right legend. Training datasets are depicted as solid lines and validation datasets are dashed lines. Losses utilized in the depicted line plot are presented to the right of each plot, see dashed rectangles. Losses #1 and #2, reconstruction and length, were trained in an alternating fashion for 20 epochs before all three losses were trained for an additional 40 epochs. C)Box plots of how reconstructed TCRs compare to the original sequence in terms of the Levenshtein distance (left) and length (right) metrics, each in AA units. Each dot represents one cross-validation iteration. A random subset of the dataset is used for training and evaluation of the metrics, which are computed on the complementary test set. D) Box plots of the Levenshtein distance comparisons of original versus reconstructed TCRs (left) and random versus reconstructed (right). P-value was computed using Mann-Whitney U test. Box plots represent median and interquartile range.

**Figure S7.**
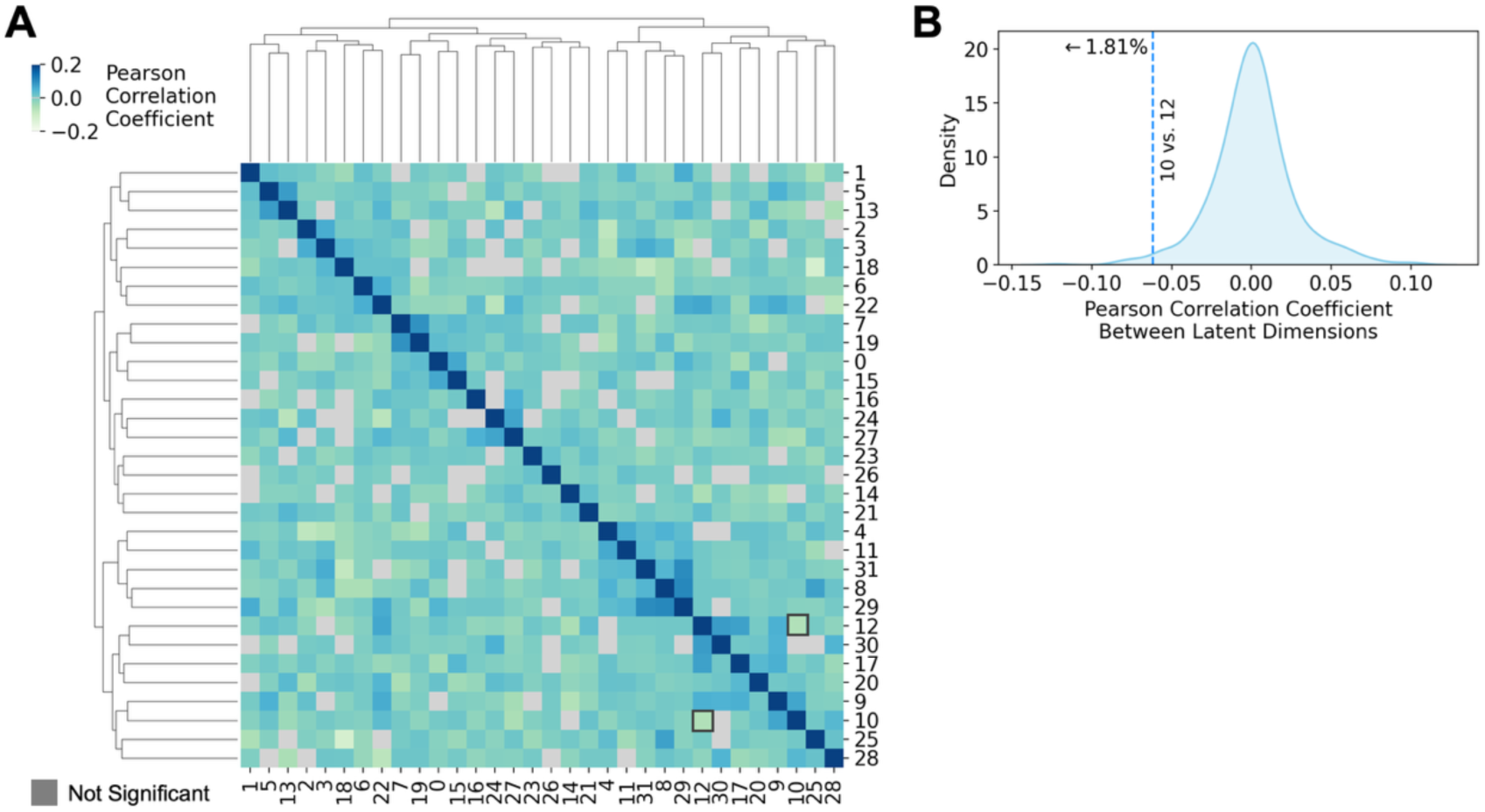
Latent dimensions comprising Tarpon TCR embeddings experience minimal correlation: A) Heatmap of the correlation coefficients between Tarpon latent dimensions. One of the most negative correlations, between latent dimensions 10 and 12, is outlined in black. Non-significant correlations are masked out in grey. B) Density plot of the correlation coefficients from panel (A). The correlation between dimensions 10 and 12 is indicated with a dashed blue line, see legend on the upper right, and the percent of correlations with at least as negative of a coefficient as the labeled one of interest is annotated on the plot. Pearson’s method was utilized for correlation.

**Figure S8.**
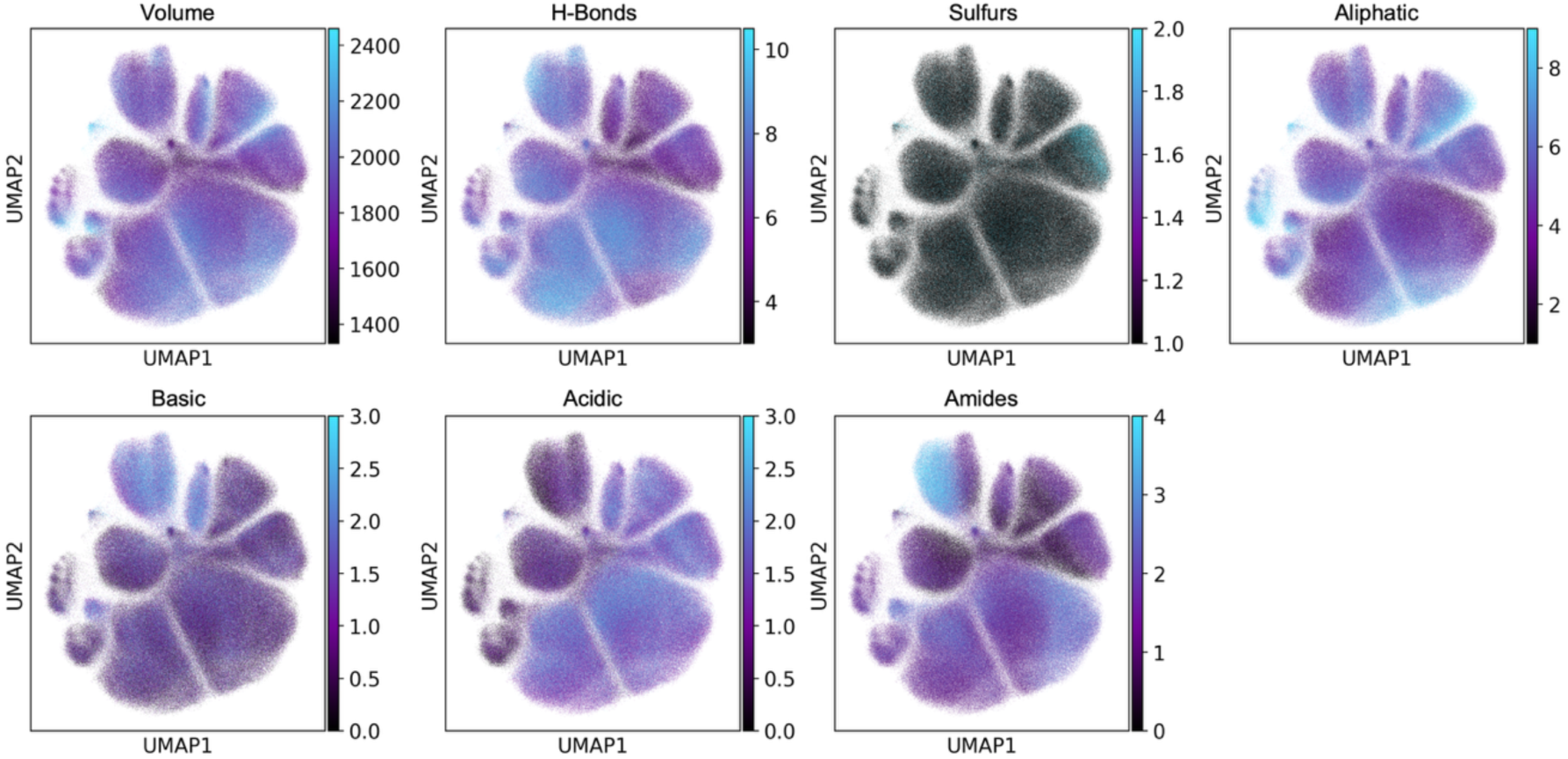
TCR physicochemical attributes are encoded in Tarpon embeddings: Selected physiochemical features of TCRs are projected onto the UMAP projection of all TCRs. The color code legend is provided at the right of each projection.

**Figure S9.**
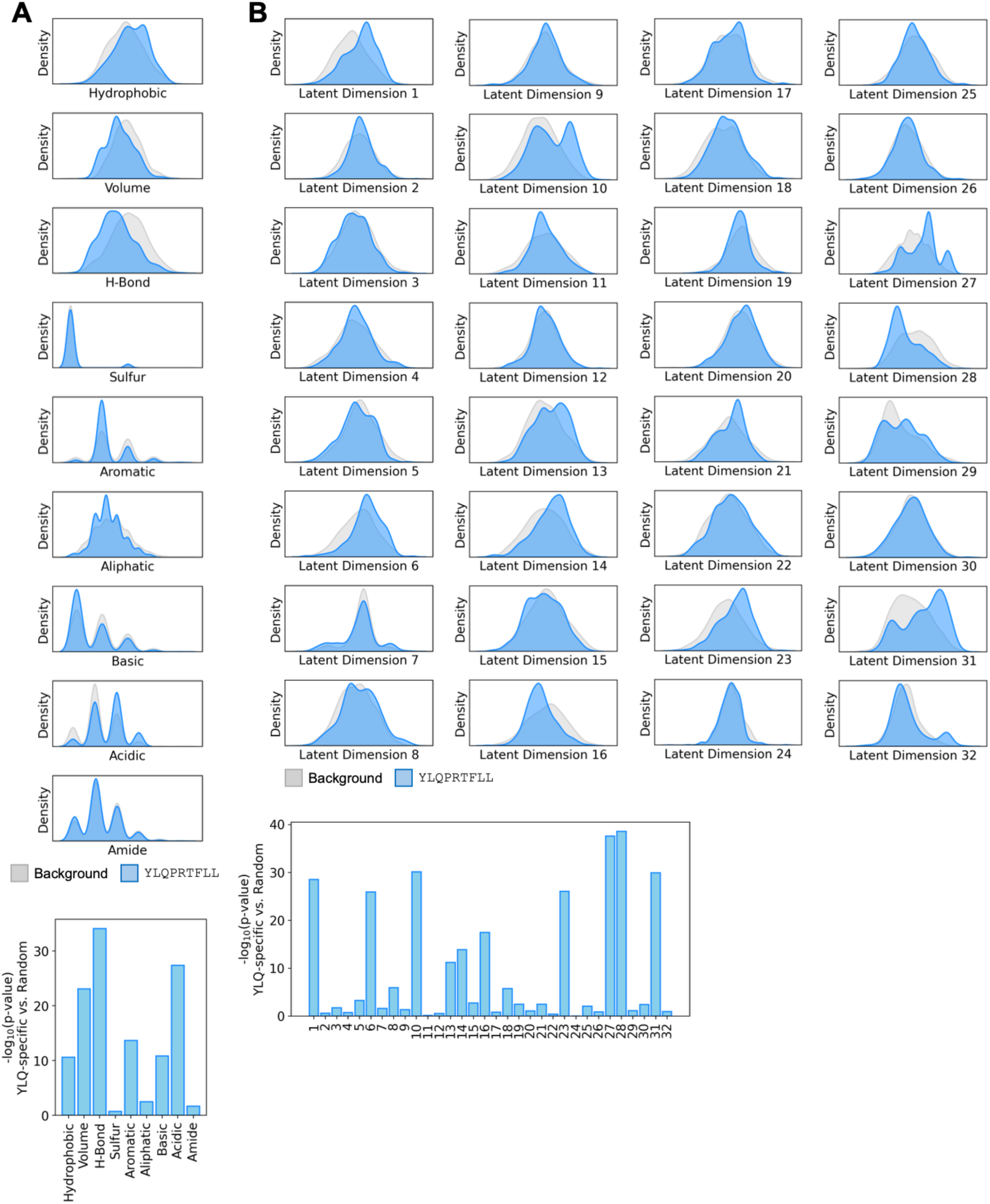
YLQPRTFLL-specific TCRs have biased physicochemical attributes and Tarpon TCR embeddings: Upper: Density plots of (A) physicochemical attributes or (B) Tarpon latent dimensions. YLQPRTFLL (YLQ) specific TCRs are in dark blue and an equal number of background, randomly selected TCRs are in light gray. Lower: Bar plots of each (A) physiochemical attribute or (B) Tarpon latent dimension. The y-axes are - log_10_(p-values) comparing the YLQ-specific TCRs against the background TCRs, using the Mann-Whitney U test.

**Figure S10.**
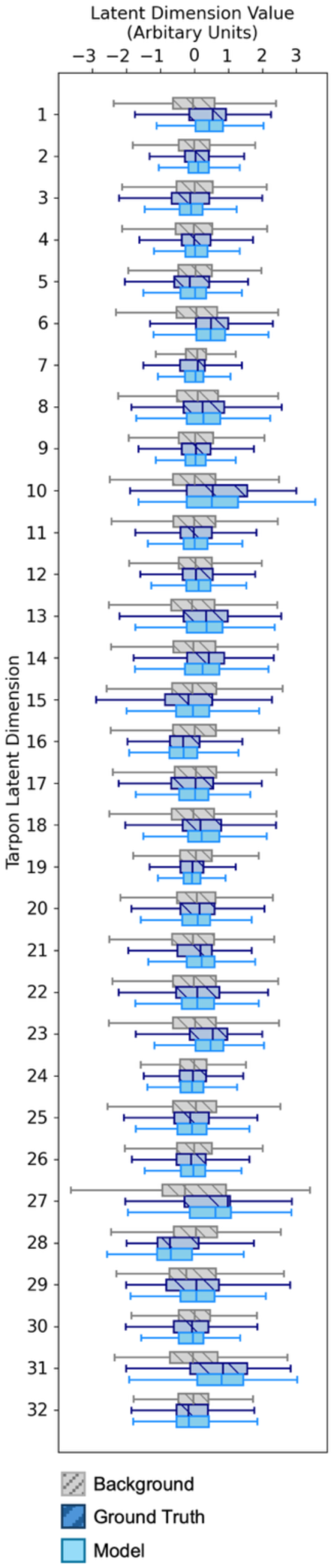
AgFlow generated YLQ-specific TCRs match the complex distribution of ground truth YLQ-Specific TCRs in Tarpon TCR space: Box plots of the Tarpon latent dimension values (x-axis) for each latent dimension (y-axis), for background random TCRs (grey), ground truth experimentally determined YLQ-specific TCRs (dark blue), and YLQ-specific TCRs generated by AgFlow (light blue). Box plots represent median and interquartile range.

**Figure S11.**
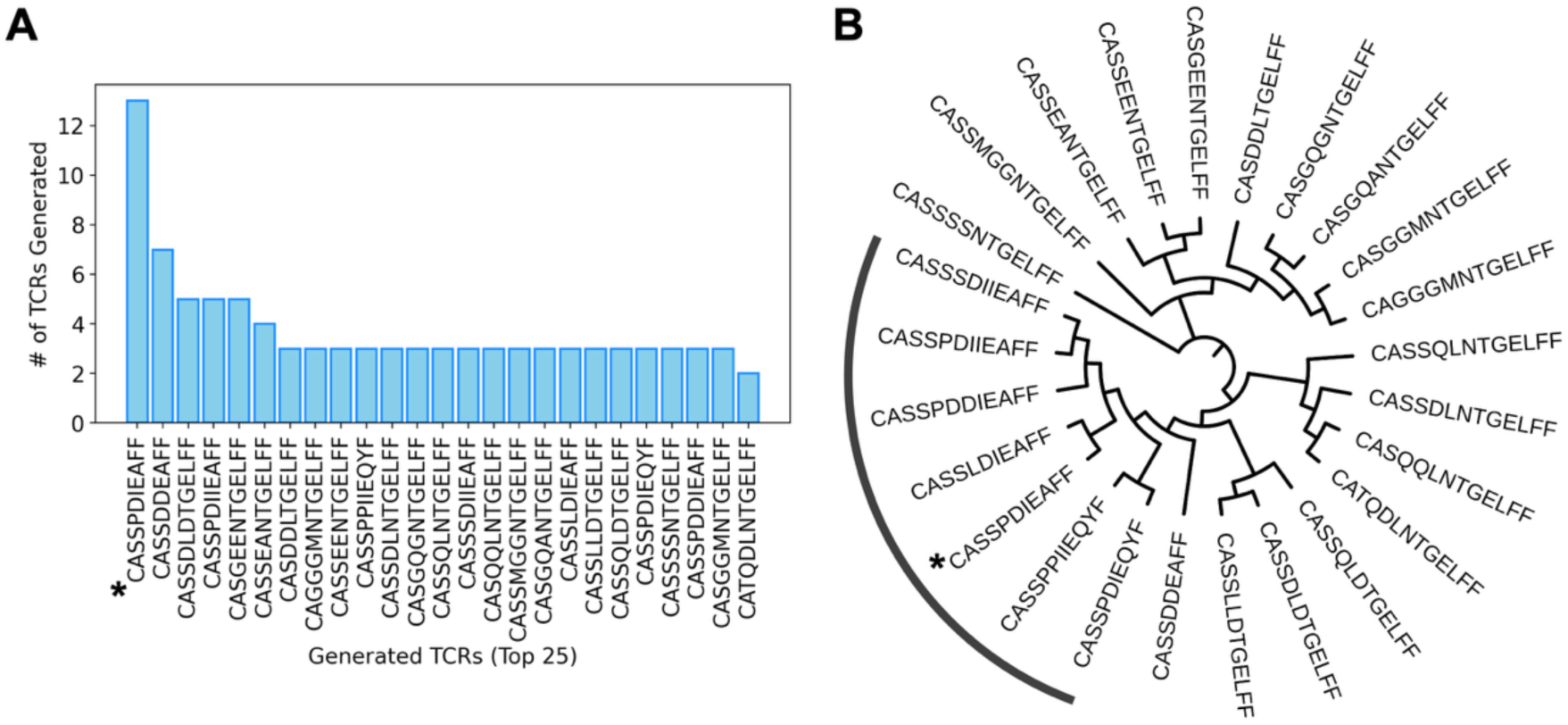
AgFlow repeatedly generates the predominant public YLQ-specific TCR: A) Bar plot of the number of TCRs generated by YLQ-specific AgFlow model. The top 25 AgFlow generated TCRs by abundance are displayed. The TCR sequence indicated with an asterisk is the predominant public YLQ-specific TCR. B) Dendrogram of TCR AA sequences from (A) with the tree computed by alignment algorithm MUSCLE. The tree is presented in branch length agnostic manner and rooted based on default parameters from MUSCLE.

**Figure S12.**
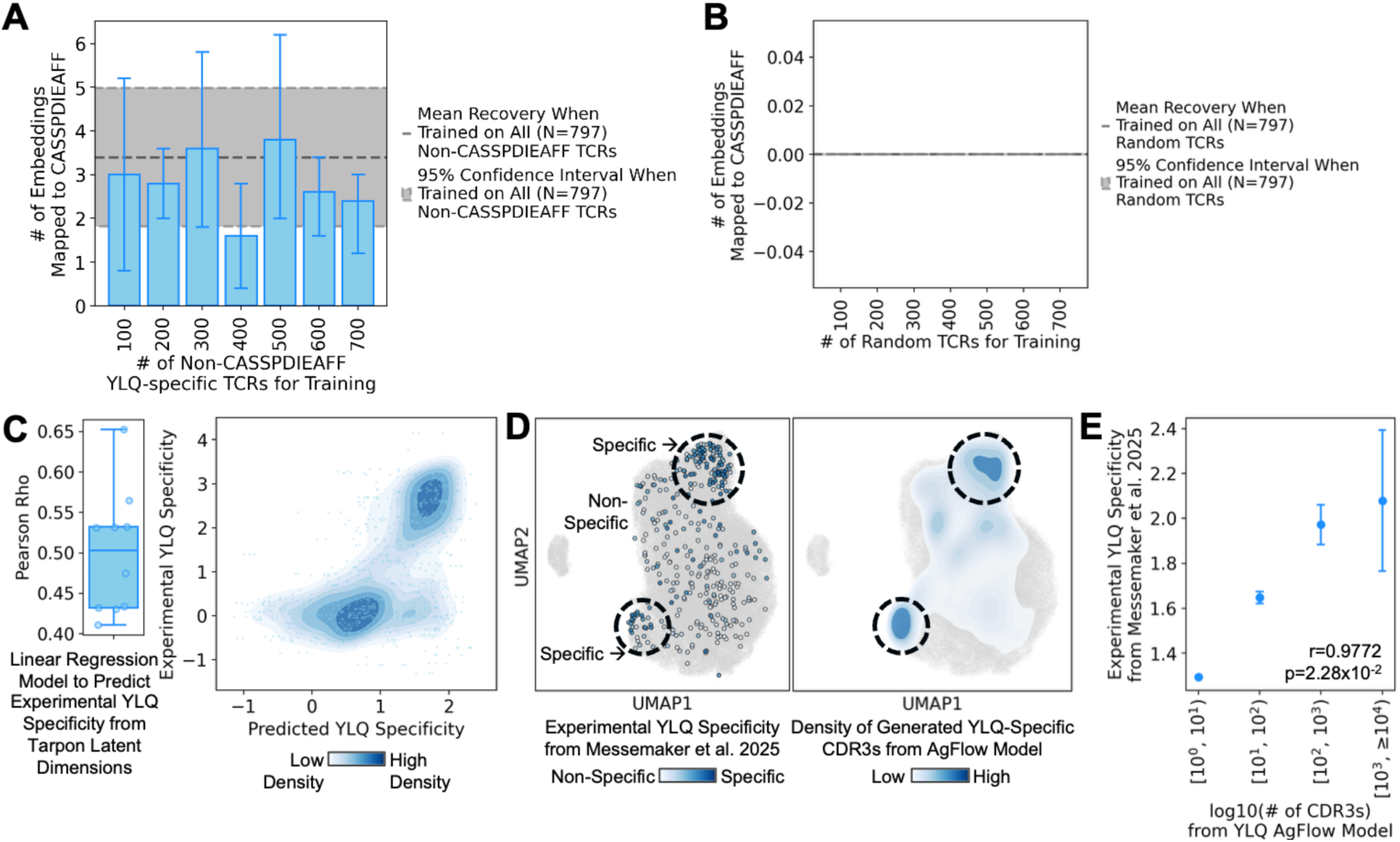
AgFlow generates the predominant public YLQ-specific TCR even when the sequence is excluded from training: A,B) Bar plot of the number of YLQ-specific TCRs utilized for training (x-axis), after the predominant public YLQ-specific TCR, CASSPDIEAFF, was removed from the training dataset against the number of AgFlow generated embeddings that mapped to this “never-seen-before” TCR (y-axis). YLQ-specific TCRs were utilized for training in (A) and random TCRs were utilized for training of the AgFlow model in (B). Bar plot heights represent arithmetic mean and error bars represent 95% confidence interval. C) Left: Pearson correlation coefficient (rho) between a given TCR’s experimental specificity to YLQ^73^ and its predicted specificity from a linear regression model trained with the given TCR’s Tarpon latent dimension values. Models were trained in a cross-validated fashion with the average rho ± 95% confidence interval as 0.50±0.05. Right: Scatterplot of predicted (x-axis) versus ground truth (y-axis) YLQ specificity scores, defined as log_2_ enrichment of a TCR’s frequency when cultured with YLQ versus irrelevant antigen expressing target cells. Density of each observation is overlaid, see color bar at the bottom. D) Scatterplot of the UMAP embedding of YLQ-specific TCRs. Sequences with experimental YLQ specificity values are presented on the left with the color indicating measured specificity by enrichment. Sequences generated by AgFlow are presented on the right with the density contour indicating areas of repeated generation. See legends on the bottom. E) Relationship between the number of times AgFlow generates a given TCR (x-axis) with the TCR’s experimentally measured YLQ specificity (y-axis). Higher y-axis values indicate increased specificity. Dots represent mean and whiskers represent 95% confidence intervals. Pearson correlation statistics are annotated on the plot.

**Figure S13.**
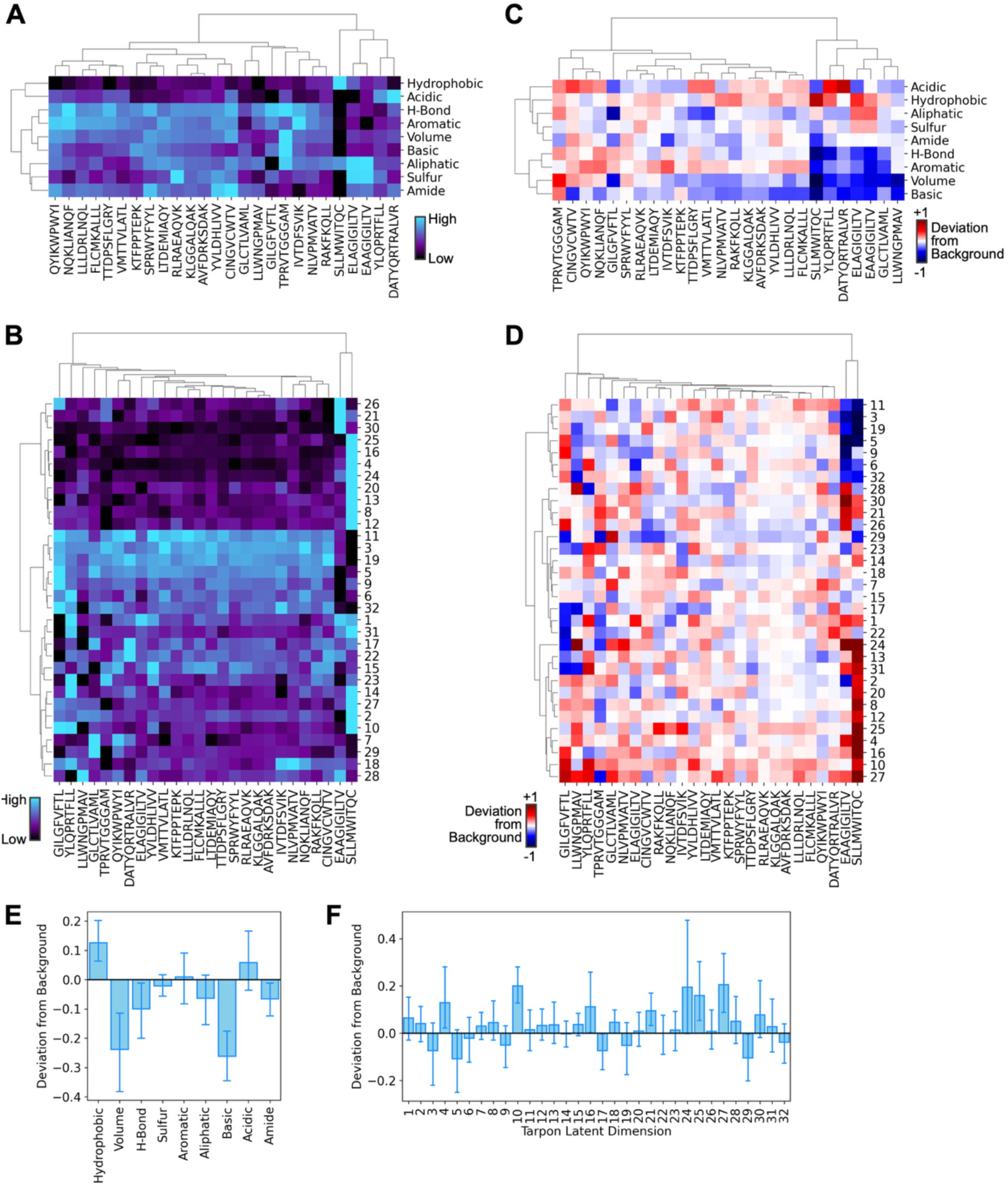
TCRs specific to known immunogenic antigens possess distinct physiochemical attributes and tarpon embeddings: A,B) Heatmap of the average (A) physicochemical attributes and (B) Tarpon latent dimensions of collections of TCRs specific to known immunogenic antigens. j Antigen sequences are provide along the x-axis. C,D) Heatmap illustrating how the average (C) physicochemical attributes and (D) Tarpon latent dimensions of collections of TCRs specific to known immunogenic antigens deviate from random TCRs, quantified by Z-score. See lower left legend E,F) Bar plots showing how the average value TCR collections specific to immunogenic antigens deviate from random TCRs by Z-score, for (E) physiochemical attributes and (F) Tarpon latent dimensions. Bar plot heights represent arithmetic mean and error bars represent 95% confidence interval.

**Figure S14.**
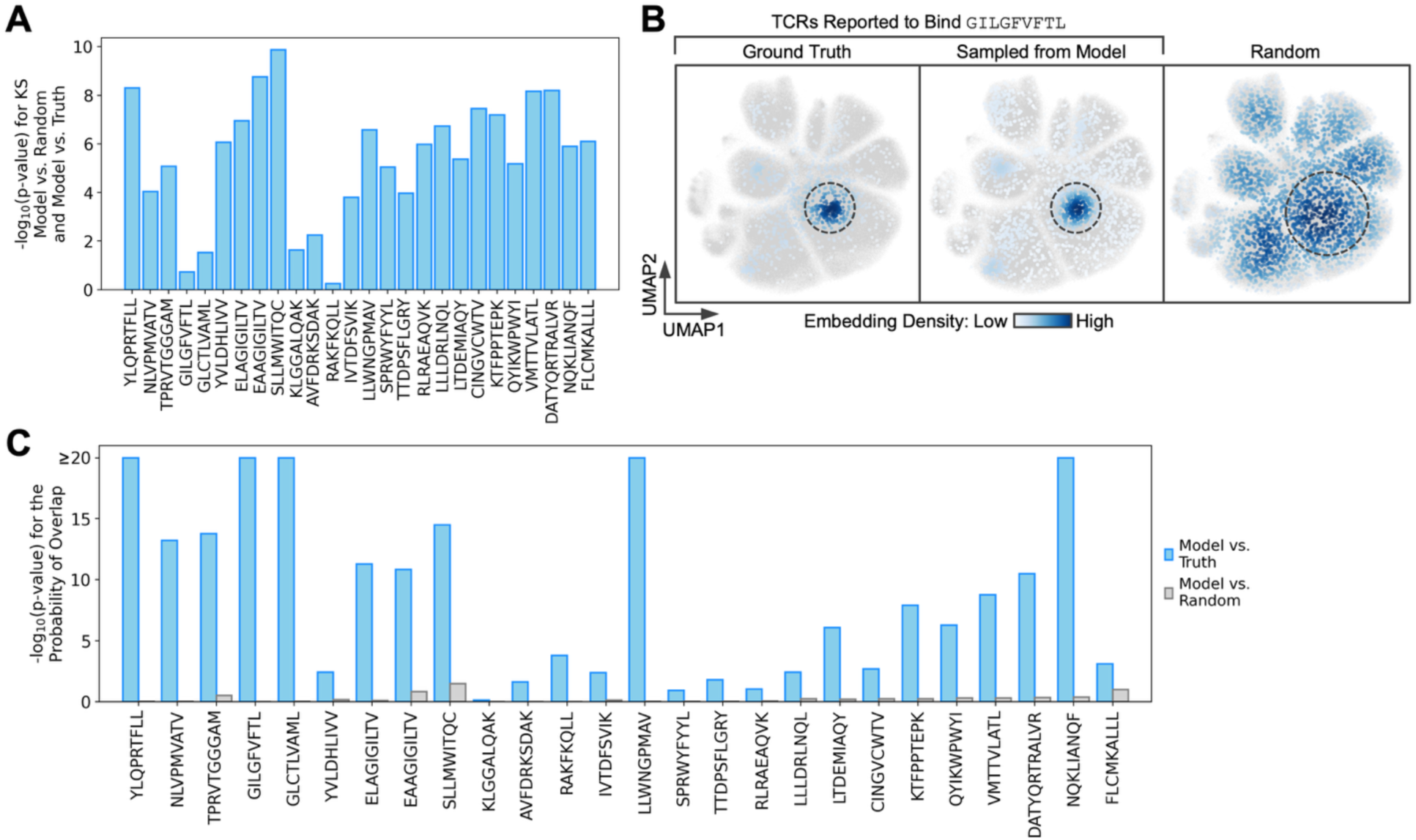
AgFlow performs well for diverse immunogenic antigens and their cognate TCRs. A) Bar plots of the difference between the Tarpon latent space distribution of AgFlow-generated TCRs for immunogenic Ags compared to experimentally derived TCRs relative to randomly selected TCRs. The y-axis is the -log_10_(p-value), from the Mann-Whitney U test, comparing the distribution differences, quantified by the Komogorov-Smirnov test, between model and background versus model and random. Lower values indicate model generated TCRs are equally similar to ground truth and random TCRs which may occur if ground truth and random TCRs possess similar Tarpon embedding distributions. Higher values are better as they indicate greater concordance with experimental data. B) GILGVFTL (influenza) specific TCRs from experimental data (ground truth, left), generated by AgFlow (middle), and random TCRs (right), as on the whole atlas Tarpon TCR embedding UMAP. Color is density (see legend). C) Bar plots of the -log_10_(p-value) that AgFlow, model, generated TCRs overlap with experimentally derived TCRs (solid blue, left of each pair), or a random set of TCRs (solid grey, right of each pair) for each immunogenic Ag. Higher values on the y-axis represent better models as they generate Ag-specific TCR repertoires with such a great overlap with the ground truth that it was unlikely to have occurred by random chance, as low p-values have high y-axis values.

**Figure S15.**
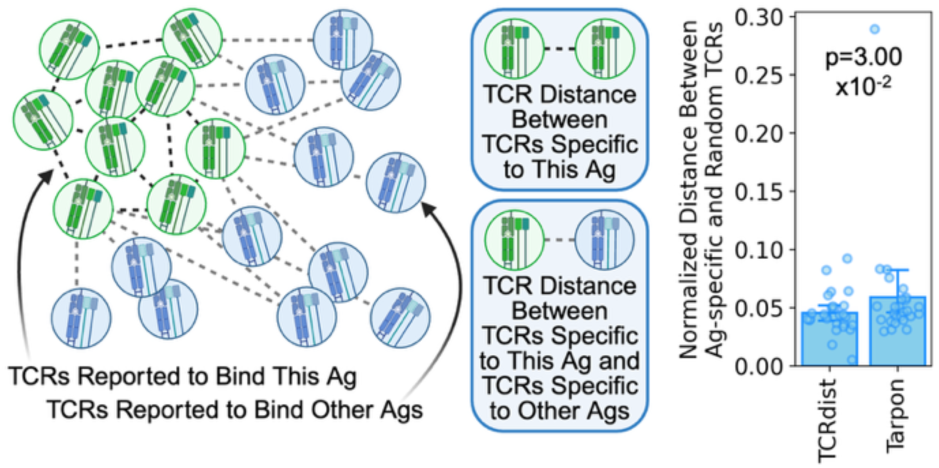
Tarpon Performs Similarly to or Better than TCRdist: Left: Illustration depicting the methodology to test how Tarpon compares to TCRdist in computing a separation metric between TCRs specific to the same antigen, relative to random, background TCRs. Right: Bar plot of the normalized distances between Ag-specific and random TCRs (y-axis), computed using TCRdist or Tarpon. The p-values were computed using the Mann-Whitney U test. Each dot represents an antigen, and many TCRs. Bar plot heights represent arithmetic mean and error bars represent 95% confidence interval. A lower height represents an improved performance.

**Figure S16.**
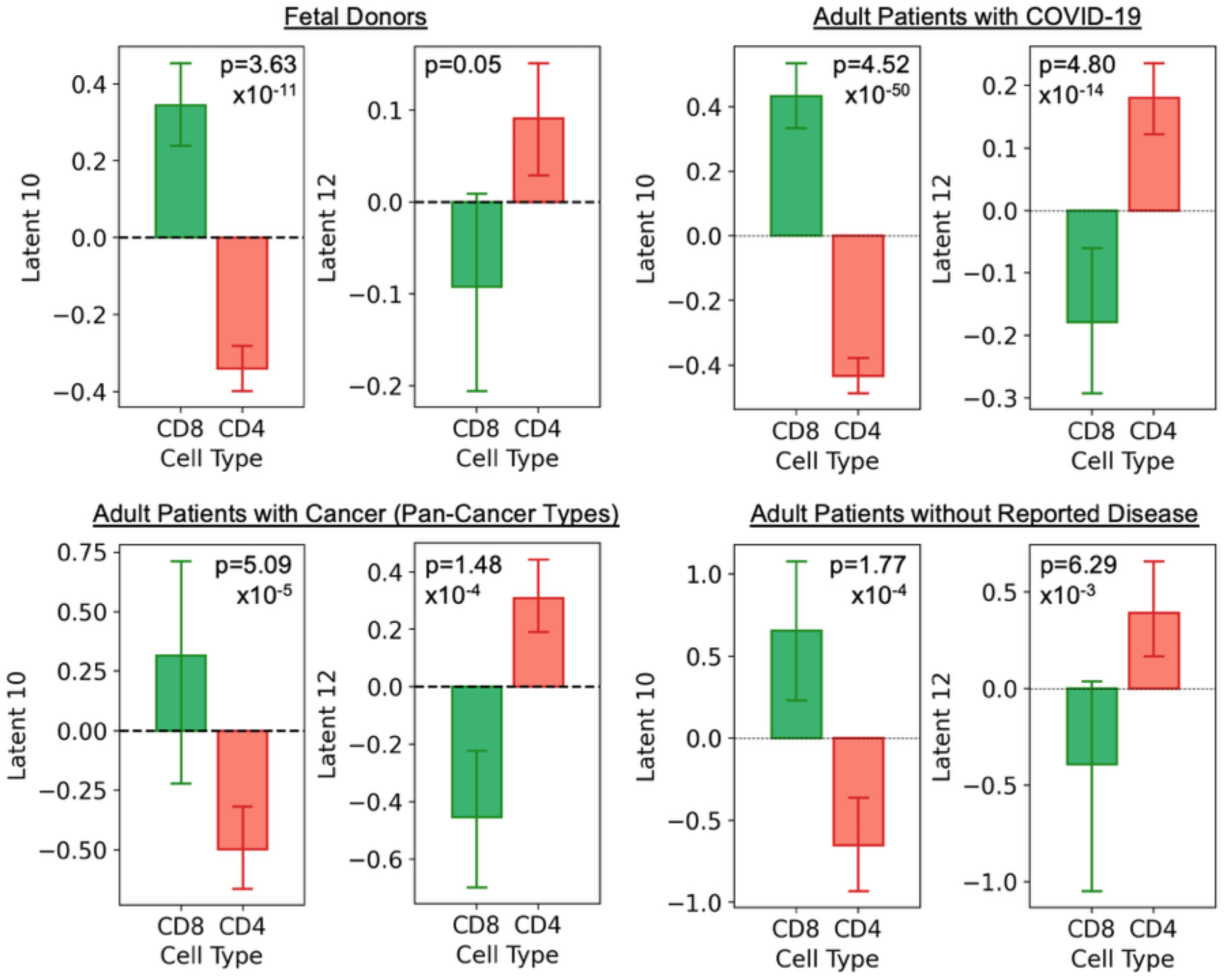
Tarpon latent dimensions 10 and 12 consistently separate single-positive CD4^+^ and CD8^+^ T cell classes: Bar plots of the latent dimension values (y-axis), for single-positive CD4^+^ and CD8^+^ TCR repertoires from donors in fetal and adult cohorts. P-values were computed using the Mann-Whitney U test. Bar plot heights represent arithmetic mean and error bars represent 95% confidence interval.

**Figure S17.**
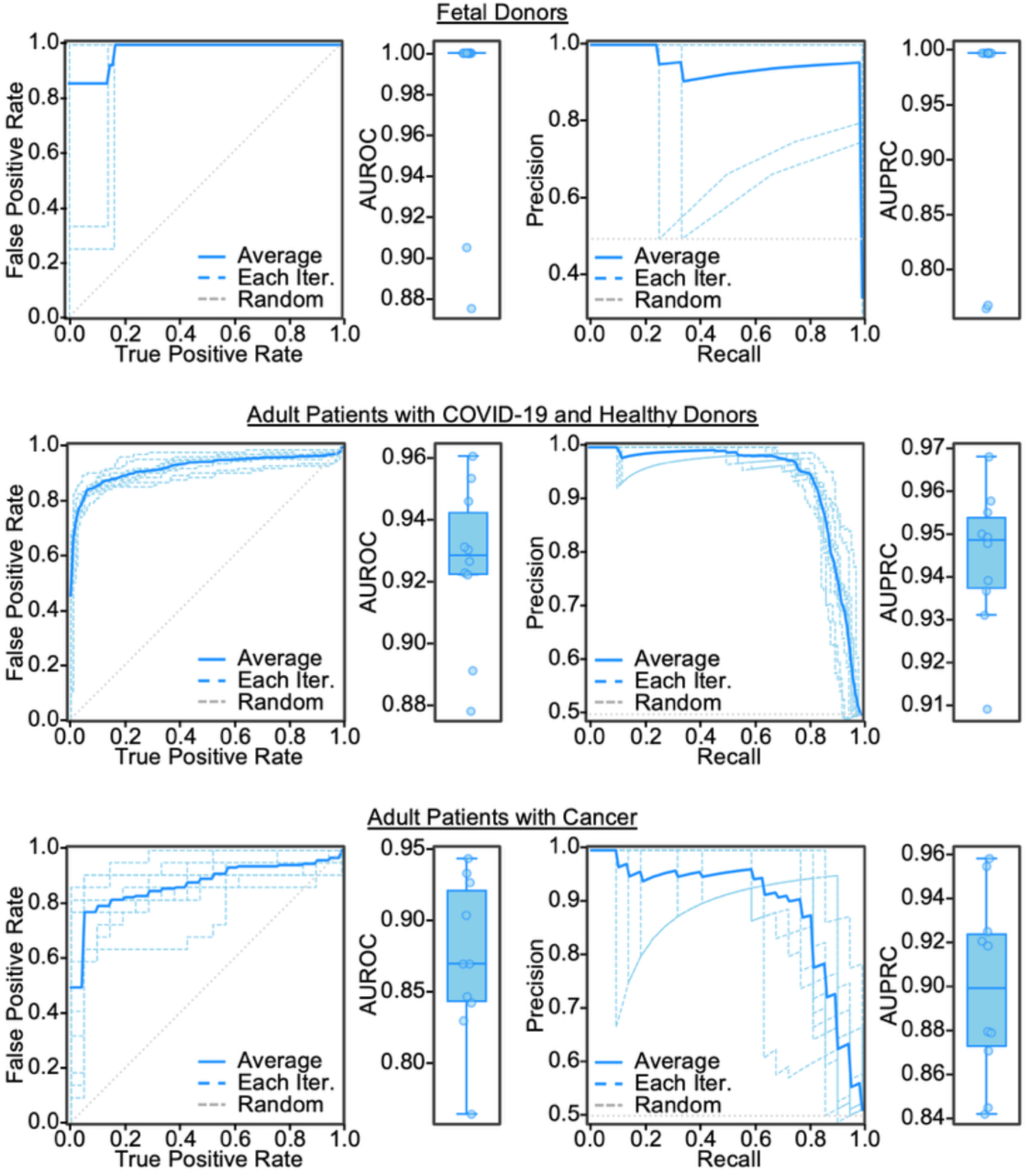
Machine learning models distinguish single-positive CD4+ and CD8+ T cell classes through their Tarpon TCR embeddings: Line plots: Receiver-operator-characteristic curves on the left and precision-recall curves on the right with false positive rate or recall on the x-axis and true positive rate or precision on the y-axis. Cross-validation iterations are represented by dashed blue lines. The average across iterations is solid blue, while random performance is a dashed grey line. Box plots: Box plots with y-axis as the area under the receiver-operator-characteristic curve (AUROC) on the left and area under the precision-recall curve (AUPRC) on the right. Models were trained and evaluated on data from fetal T cell development in upper plots, adults with COVID-19 and healthy in center plots, and adults with cancer in lower plots to distinguish single-positive CD4^+^ and CD8^+^ T cell classes. Each dot represents one cross-validation iteration. Box plots represent median and interquartile range.

**Figure S18.**
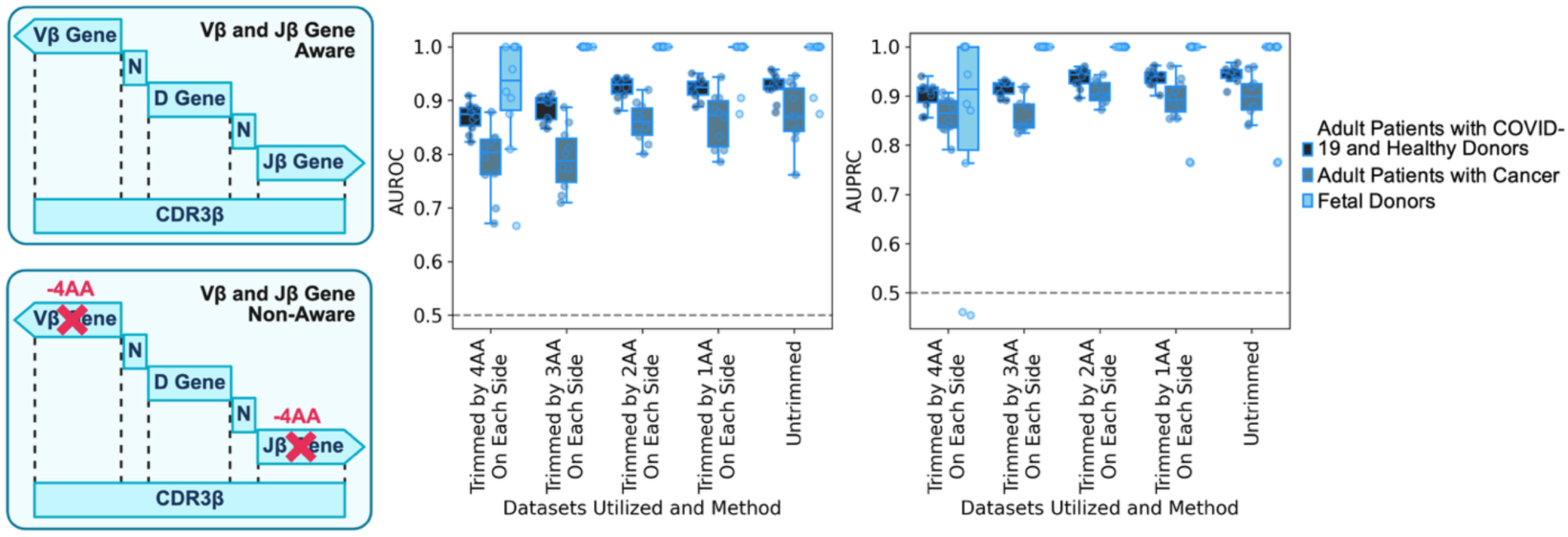
Machine learning models distinguish single-positive CD4+ and CD8+ T cell classes using Tarpon embeddings stripped of V and J gene information: Left: Illustration depicting the methodology undertaken to systematically strip away terminal amino acids of the CDR3 to remove V and J gene information and evaluate how well Tarpon embeddings can still predict the T cell class for a TCR repertoire. Right: Box plots for different iterations, whereby for each iteration, different number of amino acids are trimmed from the TCR CDR3b chains. For example, for the leftmost boxplots, 4 AAs are trimmed from both the n- and c-termini. The model performance, quantified by AUROC, is plotted on the y-axis. Each dot represents a cross-validation iteration. Color represents the cohort models used for training and evaluation (see legend on the right). Box plots represent median and interquartile range.

**Figure S19.**
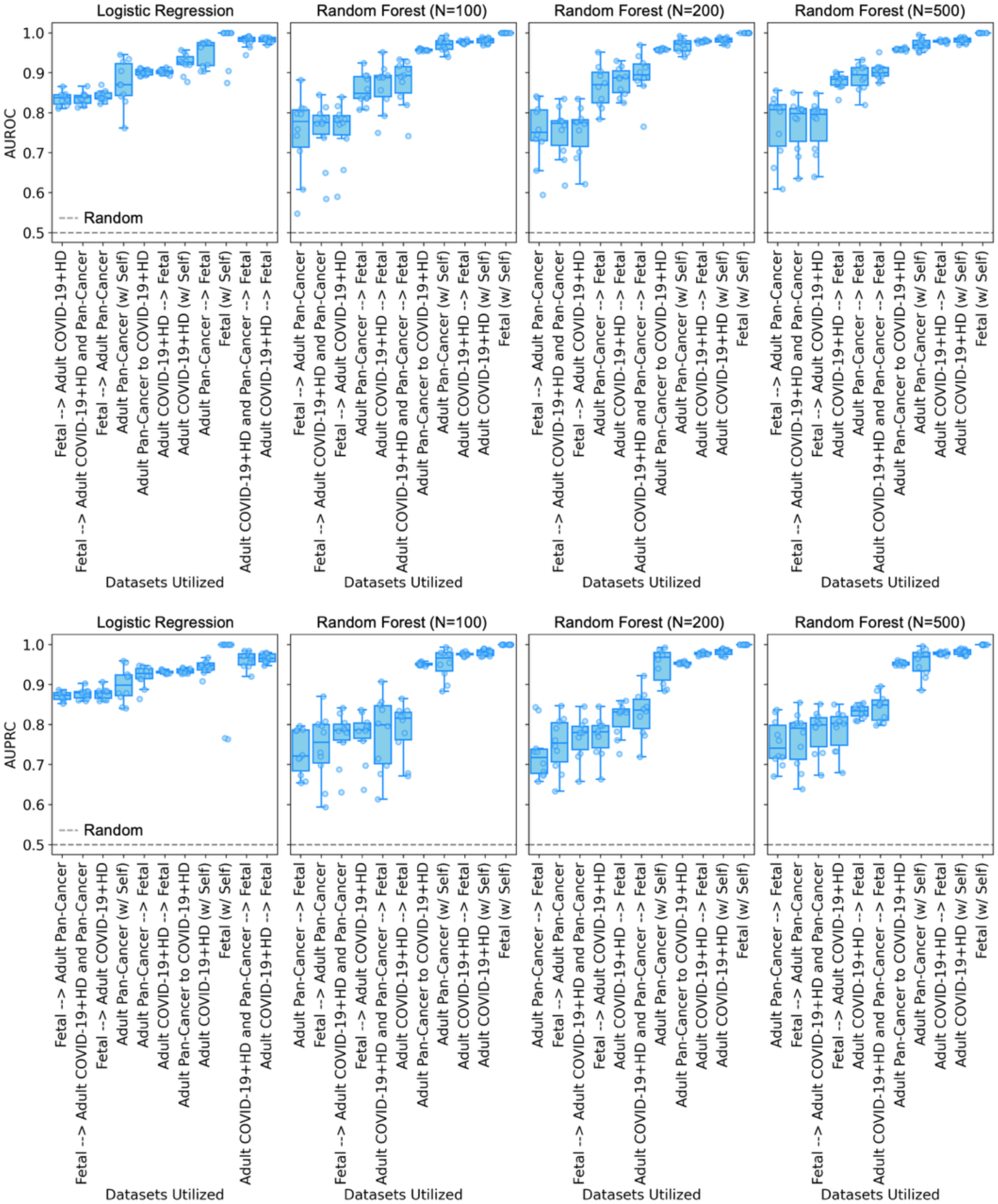
Cross-dataset mapping of Tarpon TCR fingerprints for single-positive CD4^+^ and CD8^+^ classes reveals divergent translatability: Box plots of model performance, with AUROC on the top and AUPRC on the bottom, for logistic regression and random forest machine learning models trained to distinguish single-positive CD4^+^ and CD8^+^ T cell TCR repertoires. Models are trained on each cohort and evaluated on all others, x-axis. Box plots represent median and interquartile range.

**Figure S20.**
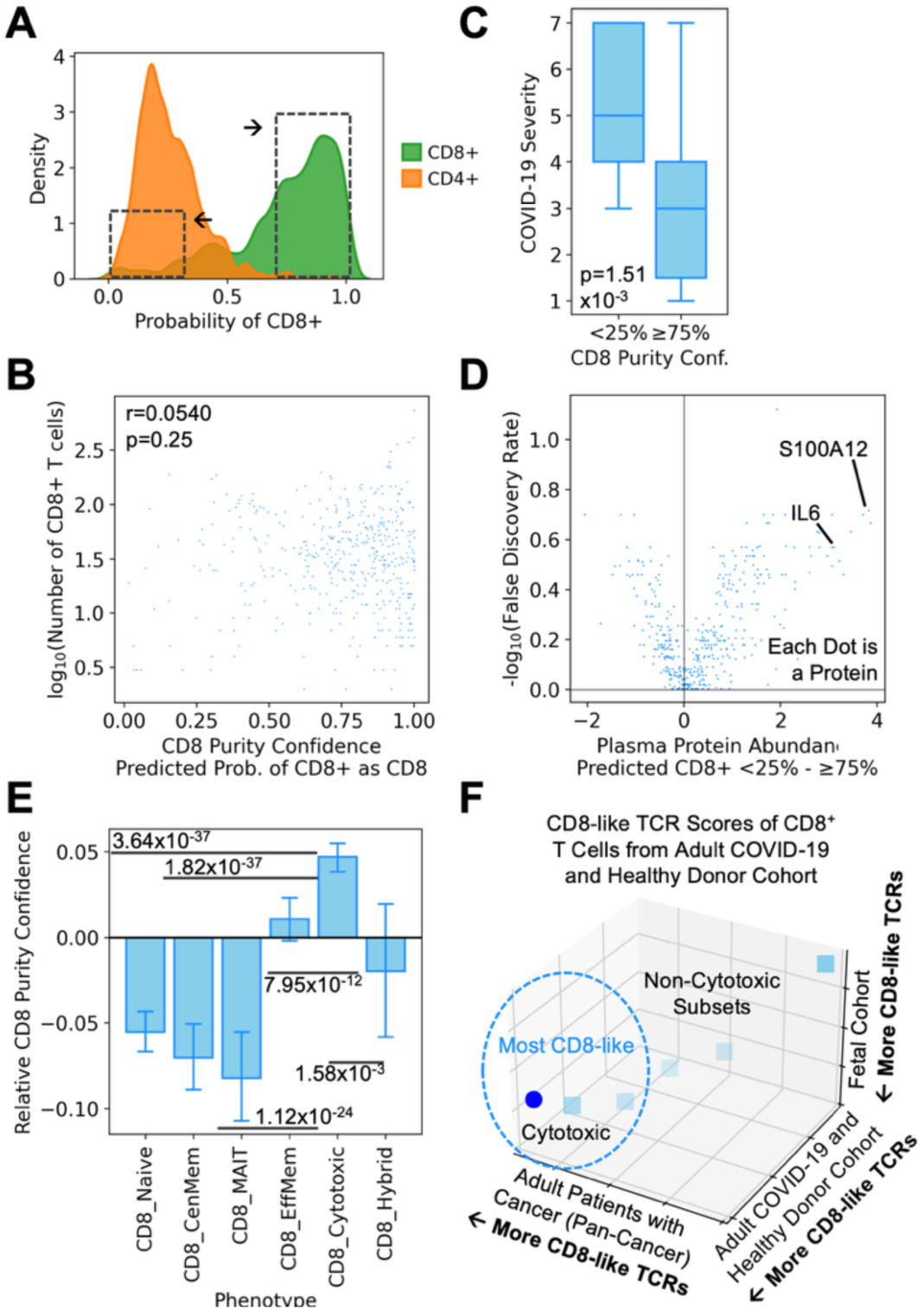
CD4-like CD8^+^ T cells associate with negative health consequences and are biased against a cytotoxic phenotype: A) Density plot of model derived CD8^+^ T cell probabilities, predicted using Tarpon TCR embeddings, for CD4^+^ (orange), and CD8^+^ (green) TCR repertoires. B) Scatterplot of model derived CD8^+^ T cell probabilities, with purity confidence (x-axis) against the numbers of CD8^+^ T cells in the given TCR repertoire. Correlation coefficient and p-value are annotated on the plot. C) Box plots of CD8^+^ TCR repertoire’s purity for individual patients, calculated from the analyses of panels (A) and (B) (x-axis), and with comparisons of COVID-19 acute severity metrics at T1 (initial diagnosis), T2 (a few days later). Disease severity was calculated using the World Health Organization (WHO) ordinal scale and the number of days a given patient spent in hospital . P-values were computed using the Mann-Whitney U test. D) Scatterplot of the differences between the plasma protein abundances (x-axis) for patients with CD8^+^ TCR repertories that are confidently predicted to have CD8-like TCRs, and of patients predicted to have CD4-like CD8^+^ T cells. The significance of these differences (y-axis) is quantified as -log_10_(False Discovery Rate, FDR). The pro-inflammatory IFNG and IL6 proteins are indicated. E) Bar plot of CD8^+^ T cell phenotype, x-axis, and, for each phenotype, a relative Z-score for CD8 purity. The scores are calculated from circulating CD8^+^ T cells, collected from patients over the course of SARS-CoV-2 infection and recovery. F) Scatterplot of panel (E) data, where each dot is a phenotype but single-positive CD4^+^ versus CD8^+^ scores are derived from TCR rulesets computed for cohorts of fetal T cell development, adults with COVID-19 and healthy donors, and adults with cancer. Cytotoxic CD8^+^ T cells are distinguished by a dark blue color and are annotated on the plot. Pearson’s method was utilized for correlation. Bar plot heights represent arithmetic mean and error bars represent 95% confidence interval. Box plots represent median and interquartile range.

**Figure S21.**
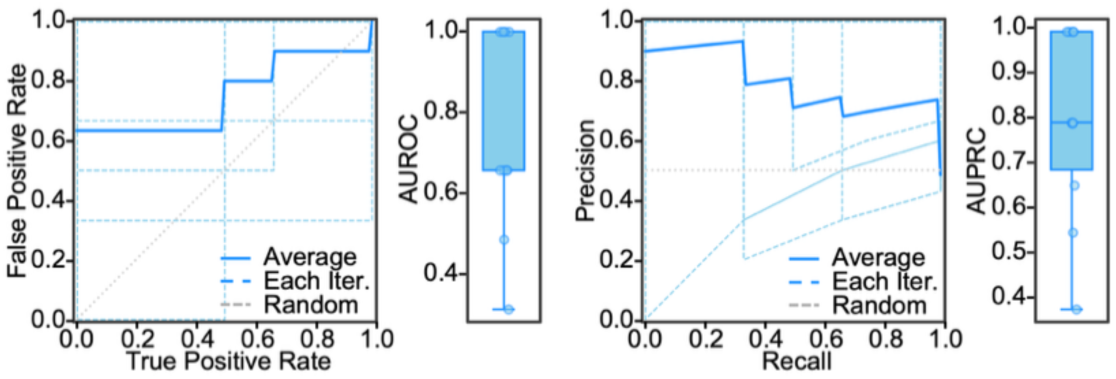
Machine learning models distinguish type I Innate T cells and conventional CD8^+^ T cells through their Tarpon TCR embeddings: Line plots: Receiver-operator-characteristic curves on the left and precision-recall curves on the right. Cross-validation iterations are represented by dashed blue lines, with the average across iterations plotted in solid blue. Random performance is presented as a dashed grey line. Box plots: Box plots of the area under the receiver-operator-characteristic curve (AUROC) (left), and area under the precision-recall curve (AUPRC) (right). Models were trained and evaluated on data from fetal T cell development to distinguish type I innate T cells from conventional CD8^+^ T cells. Each dot represents one cross-validation iteration. Box plots represent median and interquartile range.

**Figure S22.**
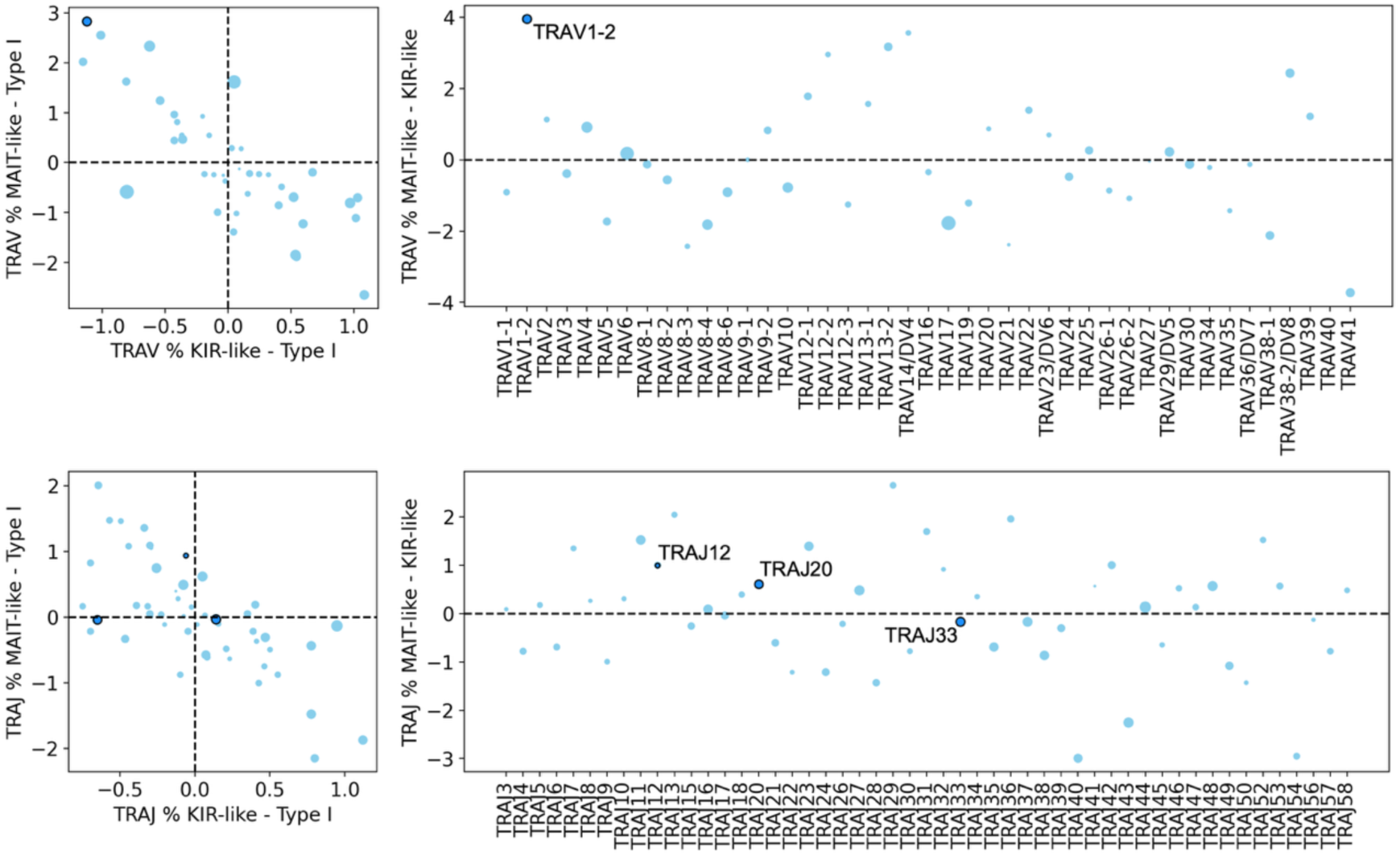
MAIT-Like Type I Innate T Cells Possess MAIT-Like V Gene Usage: Left: Scatterplots of the usage for V (upper) and J (lower) genes for MAIT and KIR like type I innate T cell subsets in comparison to the average usage of these genes in type I innate T cells. Right: Scatterplot of the difference in V (upper) and J (lower) gene usage between MAIT and KIR like type I innate T cell subsets. Genes reported to be over-represented in MAIT cells are annotated and distinguished by a dark blue color.

**Figure S23.**
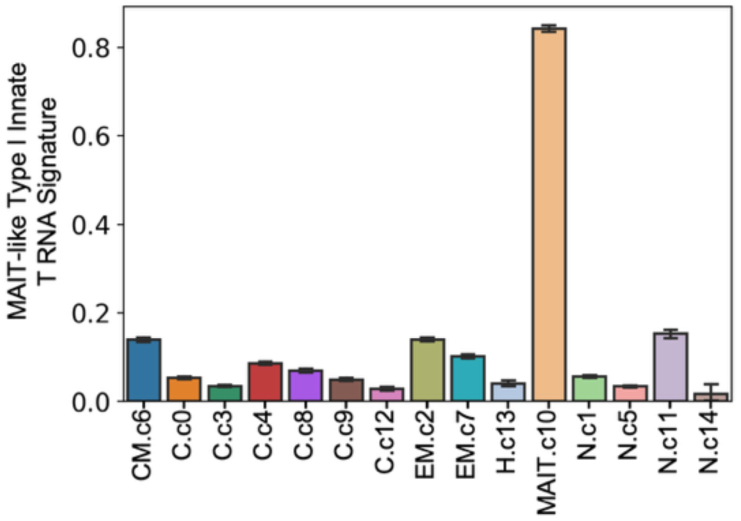
Transcriptomic Mapping of MAIT-Like Type I Innate T Cells to Adult T Cell Datasets: Celltypist mapping confidence, on a zero-to-one scale, with one being highest confidence, of MAIT-like type I innate T cells to adult T cell phenotypes. Each phenotype on the x-axis is indicated by the notation “phenotype.subcluster.. Bar plot heights represent arithmetic mean and error bars represent 95% confidence interval.

## Supplementary Tables

**Table S1:** AgFlow latent space distribution and repertoire capture metrics for all epitopes.

**Table S2:** Ag-specific TCR distance metrics from TCRdist and Tarpon.

**Table S3:** Plasma proteome differences in COVID-19 patients with atypical CD8+ TCRs.

## References

1. Zhang, N., and Bevan, M.J. (2011). CD8+ T Cells: Foot Soldiers of the Immune System. Immunity 35, 161–168. 10.1016/j.immuni.2011.07.010.

2. Borst, J., Ahrends, T., Bąbała, N., Melief, C.J.M., and Kastenmüller, W. (2018). CD4+ T cell help in cancer immunology and immunotherapy. Nat. Rev. Immunol. 18, 635–647. 10.1038/s41577-018-0044-0.

3. Toumi, R., Yuzefpolskiy, Y., Vegaraju, A., Xiao, H., Smith, K.A., Sarkar, S., and Kalia, V. (2022). Autocrine and paracrine IL-2 signals collaborate to regulate distinct phases of CD8 T cell memory. Cell Rep. 39, 110632. 10.1016/j.celrep.2022.110632.

4. Abbas, A.K. (2020). The Surprising Story of IL-2: From Experimental Models to Clinical Application. Am. J. Pathol. 190, 1776–1781. 10.1016/j.ajpath.2020.05.007.

5. Pipkin, M.E., Sacks, J.A., Cruz-Guilloty, F., Lichtenheld, M.G., Bevan, M.J., and Rao, A. (2010). Interleukin-2 and Inflammation Induce Distinct Transcriptional Programs that Promote the Differentiation of Effector Cytolytic T Cells. Immunity 32, 79–90. 10.1016/j.immuni.2009.11.012.

6. Raskov, H., Orhan, A., Christensen, J.P., and Gögenur, I. (2021). Cytotoxic CD8+ T cells in cancer and cancer immunotherapy. Br. J. Cancer 124, 359–367. 10.1038/s41416-020-01048-4.

7. Shah, K., Al-Haidari, A., Sun, J., and Kazi, J.U. (2021). T cell receptor (TCR) signaling in health and disease. Signal Transduct. Target. Ther. 6. 10.1038/s41392-021-00823-w.

8. Chen, D.G., Xie, J., Su, Y., and Heath, J.R. (2023). T cell receptor sequences are the dominant factor contributing to the phenotype of CD8+ T cells with specificities against immunogenic viral antigens. Cell Rep. 42, 113279. 10.1016/j.celrep.2023.113279.

9. Xie, J., Chen, D.G., Chour, W., Ng, R.H., Zhang, R., Yuan, D., Choi, J., McKasson, M., Troisch, P., Smith, B., et al. (2025). APMAT analysis reveals the association between CD8 T cell receptors, cognate antigen, and T cell phenotype and persistence. Nat. Commun. 16, 1402. 10.1038/s41467-025-56659-3.

10. Lagattuta, K.A., Kang, J.B., Nathan, A., Pauken, K.E., Jonsson, A.H., Rao, D.A., Sharpe, A.H., Ishigaki, K., and Raychaudhuri, S. (2022). Repertoire analyses reveal T cell antigen receptor sequence features that influence T cell fate. Nat. Immunol. 23, 446–457. 10.1038/s41590-022-01129-x.

11. Baulu, E., Gardet, C., Chuvin, N., and Depil, S. (2023). TCR-engineered T cell therapy in solid tumors: State of the art and perspectives. Sci. Adv. 9, 1–15. 10.1126/sciadv.adf3700.

12. Tsimberidou, A.-M., Van Morris, K., Vo, H.H., Eck, S., Lin, Y.-F., Rivas, J.M., and Andersson, B.S. (2021). T-cell receptor-based therapy: an innovative therapeutic approach for solid tumors. J. Hematol. Oncol. 14, 102. 10.1186/s13045-021-01115-0.

13. Oliveira, G., and Wu, C.J. (2023). Dynamics and specificities of T cells in cancer immunotherapy. Nat. Rev. Cancer 23, 295–316. 10.1038/s41568-023-00560-y.

14. Gearty, S. V., Dündar, F., Zumbo, P., Espinosa-Carrasco, G., Shakiba, M., Sanchez-Rivera, F.J., Socci, N.D., Trivedi, P., Lowe, S.W., Lauer, P., et al. (2022). An autoimmune stem-like CD8 T cell population drives type 1 diabetes. Nature 602, 156–161. 10.1038/s41586-021-04248-x.

15. Súkeníková, L., Mallone, A., Schreiner, B., Ripellino, P., Nilsson, J., Stoffel, M., Ulbrich, S.E., Sallusto, F., and Latorre, D. (2024). Autoreactive T cells target peripheral nerves in Guillain–Barré syndrome. Nature 626, 160–168. 10.1038/s41586-023-06916-6.

16. Oliveira, G., Stromhaug, K., Klaeger, S., Kula, T., Frederick, D.T., Le, P.M., Forman, J., Huang, T., Li, S., Zhang, W., et al. (2021). Phenotype, specificity and avidity of antitumour CD8+ T cells in melanoma. Nature 596, 119–125. 10.1038/s41586-021-03704-y.

17. Wang, X., Werneck, M.B.F., Wilson, B.G., Kim, H.-J., Kluk, M.J., Thom, C.S., Wischhusen, J.W., Evans, J.A., Jesneck, J.L., Nguyen, P., et al. (2011). TCR-dependent transformation of mature memory phenotype T cells in mice. J. Clin. Invest. 121, 3834–3845. 10.1172/JCI37210.

18. Han, A., Glanville, J., Hansmann, L., and Davis, M.M. (2014). Linking T-cell receptor sequence to functional phenotype at the single-cell level. Nat. Biotechnol. 32, 684–692. 10.1038/nbt.2938.

19. Zheng, L., Qin, S., Si, W., Wang, A., Xing, B., Gao, R., Ren, X., Wang, L., Wu, X., Zhang, J., et al. (2021). Pan-cancer single-cell landscape of tumor-infiltrating T cells. Science (80-.). 374. 10.1126/science.abe6474.

20. Suo, C., Dann, E., Goh, I., Jardine, L., Kleshchevnikov, V., Park, J.E., Botting, R.A., Stephenson, E., Engelbert, J., Tuong, Z.K., et al. (2022). Mapping the developing human immune system across organs. Science (80-.). *376*. 10.1126/science.abo0510.

21. Huang, H., Wang, C., Rubelt, F., Scriba, T.J., and Davis, M.M. (2020). Analyzing the Mycobacterium tuberculosis immune response by T-cell receptor clustering with GLIPH2 and genome-wide antigen screening. Nat. Biotechnol. 38, 1194– 1202. 10.1038/s41587-020-0505-4.

22. Su, Y., Yuan, D., Chen, D.G., Ng, R.H., Wang, K., Choi, J., Li, S., Hong, S., Zhang, R., Xie, J., et al. (2022). Multiple early factors anticipate post-acute COVID-19 sequelae. Cell 185, 881–895.e20. 10.1016/j.cell.2022.01.014.

23. Su, Y., Chen, D., Yuan, D., Lausted, C., Choi, J., Dai, C.L., Voillet, V., Duvvuri, V.R., Scherler, K., Troisch, P., et al. (2020). Multi-Omics Resolves a Sharp Disease-State Shift between Mild and Moderate COVID-19. Cell 183, 1479–1495.e20. 10.1016/j.cell.2020.10.037.

24. Liu, B., Hu, X., Feng, K., Gao, R., Xue, Z., Zhang, S., Zhang, Y., Corse, E., Hu, Y., Han, W., et al. (2022). Temporal single-cell tracing reveals clonal revival and expansion of precursor exhausted T cells during anti-PD-1 therapy in lung cancer 10.1038/s43018-021-00292-8.

25. Rojas, L.A., Sethna, Z., Soares, K.C., Olcese, C., Pang, N., Patterson, E., Lihm, J., Ceglia, N., Guasp, P., Chu, A., et al. (2023). Personalized RNA neoantigen vaccines stimulate T cells in pancreatic cancer. Nature 618, 144–150. 10.1038/s41586-023-06063-y.

26. Guo, X., Zhang, Y., Zheng, L., Zheng, C., Song, J., Zhang, Q., Kang, B., Liu, Z., Jin, L., Xing, R., et al. (2018). Global characterization of T cells in non-small-cell lung cancer by single-cell sequencing. Nat. Med. 24, 978–985. 10.1038/s41591-018-0045-3.

27. Barras, D., Ghisoni, E., Chiffelle, J., Orcurto, A., Dagher, J., Fahr, N., Benedetti, F., Crespo, I., Grimm, A.J., Morotti, M., et al. (2024). Response to tumor-infiltrating lymphocyte adoptive therapy is associated with preexisting CD8+ T-myeloid cell networks in melanoma. Sci. Immunol. 9, 1–16. 10.1126/sciimmunol.adg7995.

28. Minowa, T., Murata, K., Mizue, Y., Murai, A., Nakatsugawa, M., Sasaki, K., Tokita, S., Kubo, T., Kanaseki, T., Tsukahara, T., et al. (2024). Single-cell profiling of acral melanoma infiltrating lymphocytes reveals a suppressive tumor microenvironment. Sci. Transl. Med. 16, eadk8832. 10.1126/scitranslmed.adk8832.

29. Miller, A.M., Koşaloğlu-Yalçın, Z., Westernberg, L., Montero, L., Bahmanof, M., Frentzen, A., Lanka, M., Premlal, A.L.R., Seumois, G., Greenbaum, J., et al. (2024). A functional identification platform reveals frequent, spontaneous neoantigen-specific T cell responses in patients with cancer. Sci. Transl. Med. 16, 1–12. 10.1126/scitranslmed.abj9905.

30. Boland, B.S., He, Z., Tsai, M.S., Olvera, J.G., Omilusik, K.D., Duong, H.G., Kim, E.S., Limary, A.E., Jin, W., Justin Milner, J., et al. (2020). Heterogeneity and clonal relationships of adaptive immune cells in ulcerative colitis revealed by single-cell analyses. Sci. Immunol. 5. 10.1126/SCIIMMUNOL.ABB4432.

31. Naulaerts, S., Datsi, A., Borras, D.M., Martinez, A.A., Messiaen, J., Vanmeerbeek, I., Sprooten, J., Laureano, R.S., Govaerts, J., Panovska, D., et al. (2023). Multiomics and spatial mapping characterizes human CD8+ T cell states in cancer. Sci. Transl. Med. 15, 1–19. 10.1126/scitranslmed.add1016.

32. Saluzzo, S., Pandey, R.V., Gail, L.M., Dingelmaier-Hovorka, R., Kleissl, L., Shaw, L., Reininger, B., Atzmüller, D., Strobl, J., Touzeau-Römer, V., et al. (2021). Delayed antiretroviral therapy in HIV-infected individuals leads to irreversible depletion of skin- and mucosa-resident memory T cells. Immunity 54, 2842–2858.e5. 10.1016/j.immuni.2021.10.021.

33. Gao, S., Wu, Z., Arnold, B., Diamond, C., Batchu, S., Giudice, V., Alemu, L., Raffo, D.Q., Feng, X., Kajigaya, S., et al. (2022). Single-cell RNA sequencing coupled to TCR profiling of large granular lymphocyte leukemia T cells. Nat. Commun. 13. 10.1038/s41467-022-29175-x.

34. Li, H., van der Leun, A.M., Yofe, I., Lubling, Y., Gelbard-Solodkin, D., van Akkooi, A.C.J., van den Braber, M., Rozeman, E.A., Haanen, J.B.A.G., Blank, C.U., et al. (2019). Dysfunctional CD8 T Cells Form a Proliferative, Dynamically Regulated Compartment within Human Melanoma. Cell 176, 775–789.e18. 10.1016/j.cell.2018.11.043.

35. Schmidt, J., Chiffelle, J., Perez, M.A.S., Magnin, M., Bobisse, S., Arnaud, M., Genolet, R., Cesbron, J., Barras, D., Navarro Rodrigo, B., et al. (2023). Neoantigen-specific CD8 T cells with high structural avidity preferentially reside in and eliminate tumors. Nat. Commun. 14. 10.1038/s41467-023-38946-z.

36. Yost, K.E., Satpathy, A.T., Wells, D.K., Qi, Y., Wang, C., Kageyama, R., McNamara, K.L., Granja, J.M., Sarin, K.Y., Brown, R.A., et al. (2019). Clonal replacement of tumor-specific T cells following PD-1 blockade. Nat. Med. 25, 1251–1259. 10.1038/s41591-019-0522-3.

37. Caushi, J.X., Zhang, J., Ji, Z., Vaghasia, A., Zhang, B., Hsiue, E.H.C., Mog, B.J., Hou, W., Justesen, S., Blosser, R., et al. (2021). Transcriptional programs of neoantigen-specific TIL in anti-PD-1-treated lung cancers (Springer US) 10.1038/s41586-021-03752-4.

38. Xu, Y., Liu, X., Cao, X., Huang, C., Liu, E., Qian, S., Liu, X., Wu, Y., Dong, F., Qiu, C.-W., et al. (2021). Artificial intelligence: A powerful paradigm for scientific research. Innov. 2, 100179. 10.1016/j.xinn.2021.100179.

39. Gielis, S., Moris, P., Bittremieux, W., De Neuter, N., Ogunjimi, B., Laukens, K., and Meysman, P. (2020). Identification of Epitope-Specific T Cells in T-Cell Receptor Repertoires. Bioinforma. Cancer Immunother., 183–195. 10.1007/978-1-0716-0327-7_13.

40. T, R.R., Demerdash, O.N.A., and Smith, J.C. (2024). TCR-H: explainable machine learning prediction of T-cell receptor epitope binding on unseen datasets. Front. Immunol. 15, 1426173. 10.3389/fimmu.2024.1426173.

41. Lotfollahi, M., Rybakov, S., Hrovatin, K., Hediyeh-zadeh, S., Talavera-López, C., Misharin, A. V, and Theis, F.J. (2023). Biologically informed deep learning to query gene programs in single-cell atlases. Nat. Cell Biol. 25, 337–350. 10.1038/s41556-022-01072-x.

42. Kunes, R.Z., Walle, T., Land, M., Nawy, T., and Pe’er, D. (2024). Supervised discovery of interpretable gene programs from single-cell data. Nat. Biotechnol. 42, 1084–1095. 10.1038/s41587-023-01940-3.

43. Zoete, V., Irving, M., Ferber, M., Cuendet, M.A., and Michielin, O. (2013). Structure-Based, Rational Design of T Cell Receptors. Front. Immunol. 4, 268. 10.3389/fimmu.2013.00268.

44. Jones, H.F., Molvi, Z., Klatt, M.G., Dao, T., and Scheinberg, D.A. (2020). Empirical and Rational Design of T Cell Receptor-Based Immunotherapies. Front. Immunol. 11, 585385. 10.3389/fimmu.2020.585385.

45. Kingma, D.P., and Welling, M. (2019). An Introduction to Variational Autoencoders. Found. Trends® Mach. Learn. 12, 307–392. 10.1561/2200000056.

46. Lopez, R., Regier, J., Cole, M.B., Jordan, M.I., and Yosef, N. (2018). Deep generative modeling for single-cell transcriptomics. Nat. Methods 15, 1053–1058. 10.1038/s41592-018-0229-2.

47. Lotfollahi, M., Naghipourfar, M., Luecken, M.D., Khajavi, M., Büttner, M., Wagenstetter, M., Avsec, Ž., Gayoso, A., Yosef, N., Interlandi, M., et al. (2022). Mapping single-cell data to reference atlases by transfer learning. Nat. Biotechnol. 40, 121–130. 10.1038/s41587-021-01001-7.

48. Cui, H., Wang, C., Maan, H., Pang, K., Luo, F., Duan, N., and Wang, B. (2024). scGPT: toward building a foundation model for single-cell multi-omics using generative AI. Nat. Methods 21, 1470–1480. 10.1038/s41592-024-02201-0.

49. Wong, F., Zheng, E.J., Valeri, J.A., Donghia, N.M., Anahtar, M.N., Omori, S., Li, A., Cubillos-Ruiz, A., Krishnan, A., Jin, W., et al. (2023). Discovery of a structural class of antibiotics with explainable deep learning (Springer US) 10.1038/s41586-023-06887-8.

50. M. Bran, A., Cox, S., Schilter, O., Baldassari, C., White, A.D., and Schwaller, P. (2024). Augmenting large language models with chemistry tools. Nat. Mach. Intell. 6, 525–535. 10.1038/s42256-024-00832-8.

51. Boiko, D.A., MacKnight, R., Kline, B., and Gomes, G. (2023). Autonomous chemical research with large language models. Nature 624, 570–578. 10.1038/s41586-023-06792-0.

52. Wang, J., Lisanza, S., Juergens, D., Tischer, D., Watson, J.L., Castro, K.M., Ragotte, R., Saragovi, A., Milles, L.F., Baek, M., et al. (2022). Scaffolding protein functional sites using deep learning. Science (80-.). 377, 387–394. 10.1126/science.abn2100.

53. Rosati, E., Dowds, C.M., Liaskou, E., Henriksen, E.K.K., Karlsen, T.H., and Franke, A. (2017). Overview of methodologies for T-cell receptor repertoire analysis. BMC Biotechnol. 17, 1–16. 10.1186/s12896-017-0379-9.

54. Turner, S.J., Doherty, P.C., McCluskey, J., and Rossjohn, J. (2006). Structural determinants of T-cell receptor bias in immunity. Nat. Rev. Immunol. 6, 883–894. 10.1038/nri1977.

55. Goncharov, M., Bagaev, D., Shcherbinin, D., Zvyagin, I., Bolotin, D., Thomas, P.G., Minervina, A.A., Pogorelyy, M. V., Ladell, K., McLaren, J.E., et al. (2022). VDJdb in the pandemic era: a compendium of T cell receptors specific for SARS-CoV-2. Nat. Methods 19, 1017–1019. 10.1038/s41592-022-01578-0.

56. Tickotsky, N., Sagiv, T., Prilusky, J., Shifrut, E., and Friedman, N. (2017). McPAS-TCR: A manually curated catalogue of pathology-associated T cell receptor sequences. Bioinformatics 33, 2924–2929. 10.1093/bioinformatics/btx286.

57. Vita, R., Mahajan, S., Overton, J.A., Dhanda, S.K., Martini, S., Cantrell, J.R., Wheeler, D.K., Sette, A., and Peters, B. (2019). The Immune Epitope Database (IEDB): 2018 update. Nucleic Acids Res. 47, D339–D343. 10.1093/nar/gky1006.

58. Zaslavsky, M.E., Craig, E., Michuda, J.K., Sehgal, N., Ram-Mohan, N., Lee, J.-Y., Nguyen, K.D., Hoh, R.A., Pham, T.D., Röltgen, K., et al. (2025). Disease diagnostics using machine learning of B cell and T cell receptor sequences. Science (80-.). 387, eadp2407. 10.1126/science.adp2407.

59. Bachmann, M. (2025). Levenshtein.

60. Gupta, A., Wefers, Z., Kahnert, K., Hansen, J.N., Leineweber, W., Cesnik, A., Lu, D., Axelsson, U., Ballllosera Navarro, F., Karaletsos, T., et al. (2024). SubCell: Vision foundation models for microscopy capture single-cell biology. bioRxiv, 2024.12.06.627299. 10.1101/2024.12.06.627299.

61. Dalla-Torre, H., Gonzalez, L., Mendoza-Revilla, J., Lopez Carranza, N., Grzywaczewski, A.H., Oteri, F., Dallago, C., Trop, E., de Almeida, B.P., Sirelkhatim, H., et al. (2025). Nucleotide Transformer: building and evaluating robust foundation models for human genomics. Nat. Methods 22, 287–297. 10.1038/s41592-024-02523-z.

62. Sethna, Z., Elhanati, Y., Callan Jr, C.G., Walczak, A.M., and Mora, T. (2019). OLGA: fast computation of generation probabilities of B- and T-cell receptor amino acid sequences and motifs. Bioinformatics 35, 2974–2981. 10.1093/bioinformatics/btz035.

63. McInnes, L., Healy, J., and Melville, J. (2018). UMAP: Uniform Manifold Approximation and Projection for Dimension Reduction.

64. Traag, V.A., Waltman, L., and van Eck, N.J. (2019). From Louvain to Leiden: guaranteeing well-connected communities. Sci. Rep. 9. 10.1038/s41598-019-41695-z.

65. Hudson, D., Fernandes, R.A., Basham, M., Ogg, G., and Koohy, H. (2023). Can we predict T cell specificity with digital biology and machine learning? Nat. Rev. Immunol. 23, 511–521. 10.1038/s41577-023-00835-3.

66. Moris, P., De Pauw, J., Postovskaya, A., Gielis, S., De Neuter, N., Bittremieux, W., Ogunjimi, B., Laukens, K., and Meysman, P. (2021). Current challenges for unseen-epitope TCR interaction prediction and a new perspective derived from image classification. Brief. Bioinform. 22, 1–12. 10.1093/bib/bbaa318.

67. Yang, M., Huang, Z.-A., Zhou, W., Ji, J., Zhang, J., He, S., and Zhu, Z. (2023). MIX-TPI: a flexible prediction framework for TCR–pMHC interactions based on multimodal representations. Bioinformatics 39, btad475. 10.1093/bioinformatics/btad475.

68. Szeto, C., Nguyen, A.T., Lobos, C.A., Chatzileontiadou, D.S.M., Jayasinghe, D., Grant, E.J., Riboldi-Tunnicliffe, A., Smith, C., and Gras, S. (2021). Molecular Basis of a Dominant SARS-CoV-2 Spike-Derived Epitope Presented by HLA-A*02:01 Recognised by a Public TCR. Cells 10. 10.3390/cells10102646.

69. Kobyzev, I., Prince, S.J.D., and Brubaker, M.A. (2021). Normalizing Flows: An Introduction and Review of Current Methods. IEEE Trans. Pattern Anal. Mach. Intell. 43, 3964–3979. 10.1109/tpami.2020.2992934.

70. Papamakarios, G., Nalisnick, E., Rezende, D.J., Mohamed, S., and Lakshminarayanan, B. (2021). Normalizing Flows for Probabilistic Modeling and Inference. arXiv.

71. Armistead, B., Jiang, Y., Carlson, M., Ford, E.S., Jani, S., Houck, J., Wu, X., Jing, L., Pecor, T., Kachikis, A., et al. (2023). Spike-specific T cells are enriched in breastmilk following SARS-CoV-2 mRNA vaccination. Mucosal Immunol. 16, 39–49. 10.1016/j.mucimm.2023.01.003.

72. Bieberich, F., Vazquez-Lombardi, R., Yermanos, A., Ehling, R.A., Mason, D.M., Wagner, B., Kapetanovic, E., Di Roberto, R.B., Weber, C.R., Savic, M., et al. (2021). A Single-Cell Atlas of Lymphocyte Adaptive Immune Repertoires and Transcriptomes Reveals Age-Related Differences in Convalescent COVID-19 Patients. Front. Immunol. 12, 701085. 10.3389/fimmu.2021.701085.

73. Messemaker, M., Kwee, B.P.Y., Moravec, Ž., Álvarez-Salmoral, D., Urbanus, J., de Paauw, S., Geerligs, J., Voogd, R., Morris, B., Guislain, A., et al. (2025). A functionally validated TCR-pMHC database for TCR specificity model development. bioRxiv, 2025.04.28.651095. 10.1101/2025.04.28.651095.

74. Leary, A.Y., Scott, D., Gupta, N.T., Waite, J.C., Skokos, D., Atwal, G.S., and Hawkins, P.G. (2024). Designing meaningful continuous representations of T cell receptor sequences with deep generative models. Nat. Commun. 15, 4271. 10.1038/s41467-024-48198-0.

75. Suo, C., Polanski, K., Dann, E., Lindeboom, R.G.H., Vilarrasa-Blasi, R., Vento-Tormo, R., Haniffa, M., Meyer, K.B., Dratva, L.M., Tuong, Z.K., et al. (2024). Dandelion uses the single-cell adaptive immune receptor repertoire to explore lymphocyte developmental origins. Nat. Biotechnol. 42, 40–51. 10.1038/s41587-023-01734-7.

76. Lindeboom, R.G.H., Worlock, K.B., Dratva, L.M., Yoshida, M., Scobie, D., Wagstaffe, H.R., Richardson, L., Wilbrey-Clark, A., Barnes, J.L., Kretschmer, L., et al. (2024). Human SARS-CoV-2 challenge uncovers local and systemic response dynamics. Nature 631, 189–198. 10.1038/s41586-024-07575-x.

77. Drost, F., An, Y., Bonafonte-Pardàs, I., Dratva, L.M., Lindeboom, R.G.H., Haniffa, M., Teichmann, S.A., Theis, F., Lotfollahi, M., and Schubert, B. (2024). Multi-modal generative modeling for joint analysis of single-cell T cell receptor and gene expression data. Nat. Commun. 15, 5577. 10.1038/s41467-024-49806-9.

78. Mayer-Blackwell, K., Fiore-Gartland, A., and Thomas, P.G. (2022). Flexible Distance-Based TCR Analysis in Python with tcrdist3. Methods Mol. Biol., 309–366. 10.1007/978-1-0716-2712-9_16.

79. Mayer-Blackwell, K., Schattgen, S., Cohen-Lavi, L., Crawford, J.C., Souquette, A., Gaevert, J.A., Hertz, T., Thomas, P.G., Bradley, P., and Fiore-Gartland, A. (2021). TCR meta-clonotypes for biomarker discovery with tcrdist3 enabled identification of public, HLA-restricted clusters of SARS-CoV-2 TCRs. Elife 10, e68605. 10.7554/eLife.68605.

80. Hernández-Hoyos, G., Anderson, M.K., Wang, C., Rothenberg, E. V, and Alberola-Ila, J. (2003). GATA-3 Expression Is Controlled by TCR Signals and Regulates CD4/CD8 Differentiation. Immunity 19, 83–94. 10.1016/S1074-7613(03)00176-6.

81. Tubo, N.J., and Jenkins, M.K. (2014). TCR signal quantity and quality in CD4+ T cell differentiation. Trends Immunol. 35, 591–596. 10.1016/j.it.2014.09.008.

82. Cox, D.R., and Snell, E.J. (1989). Analysis of Binary Data 2nd ed. (Chapman & Hall/CRC).

83. Davis, M.M., and Bjorkman, P.J. (1988). T-cell antigen receptor genes and T-cell recognition. Nature 334, 395–402. 10.1038/334395a0.

84. Pellicci, D.G., Koay, H.F., and Berzins, S.P. (2020). Thymic development of unconventional T cells: how NKT cells, MAIT cells and γδ T cells emerge. Nat. Rev. Immunol. 20, 756–770. 10.1038/s41577-020-0345-y.

85. Godfrey, D.I., Koay, H.F., McCluskey, J., and Gherardin, N.A. (2019). The biology and functional importance of MAIT cells. Nat. Immunol. 20, 1110–1128. 10.1038/s41590-019-0444-8.

86. Li, H., Ye, C., Ji, G., and Han, J. (2012). Determinants of public T cell responses. Cell Res. 22, 33–42. 10.1038/cr.2012.1.

87. Shomuradova, A.S., Vagida, M.S., Sheetikov, S.A., Zornikova, K. V., Kiryukhin, D., Titov, A., Peshkova, I.O., Khmelevskaya, A., Dianov, D. V., Malasheva, M., et al. (2020). SARS-CoV-2 Epitopes Are Recognized by a Public and Diverse Repertoire of Human T Cell Receptors. Immunity 53, 1245–1257.e5. 10.1016/j.immuni.2020.11.004.

88. Sidhom, J.-W., Larman, H.B., Pardoll, D.M., and Baras, A.S. (2021). DeepTCR is a deep learning framework for revealing sequence concepts within T-cell repertoires. Nat. Commun. 12, 1605. 10.1038/s41467-021-21879-w.

89. Xu, A.M., Chour, W., DeLucia, D.C., Su, Y., Pavlovitch-Bedzyk, A.J., Ng, R., Rasheed, Y., Davis, M.M., Lee, J.K., and Heath, J.R. (2023). Entropic analysis of antigen-specific CDR3 domains identifies essential binding motifs shared by CDR3s with different antigen specificities. Cell Syst. 14, 273–284.e5. 10.1016/j.cels.2023.03.001.

90. Stadinski, B.D., Shekhar, K., Gómez-Touriño, I., Jung, J., Sasaki, K., Sewell, A.K., Peakman, M., Chakraborty, A.K., and Huseby, E.S. (2016). Hydrophobic CDR3 residues promote the development of self-reactive T cells. Nat. Immunol. 17, 946– 955. 10.1038/ni.3491.

91. Ma, K.-Y., Schonnesen, A.A., He, C., Xia, A.Y., Sun, E., Chen, E., Sebastian, K.R., Guo, Y.-W., Balderas, R., Kulkarni-Date, M., et al. (2021). High-throughput and high-dimensional single-cell analysis of antigen-specific CD8+ T cells. Nat. Immunol. 22, 1590–1598. 10.1038/s41590-021-01073-2.

92. Sharon, E., Sibener, L. V, Battle, A., Fraser, H.B., Garcia, K.C., and Pritchard, J.K. (2016). Genetic variation in MHC proteins is associated with T cell receptor expression biases. Nat. Genet. 48, 995–1002. 10.1038/ng.3625.

93. McMaster, B., Thorpe, C.J., Rossjohn, J., Deane, C.M., and Koohy, H. (2024). Quantifying conformational changes in the TCR:pMHC-I binding interface. Front. Immunol. 15, 1491656. 10.3389/fimmu.2024.1491656.

94. Manso, T., Folch, G., Giudicelli, V., Jabado-Michaloud, J., Kushwaha, A., Nguefack Ngoune, V., Georga, M., Papadaki, A., Debbagh, C., Pégorier, P., et al. (2022). IMGT® databases, related tools and web resources through three main axes of research and development. Nucleic Acids Res. 50, D1262–D1272. 10.1093/nar/gkab1136.

95. Chen, S.-Y., Yue, T., Lei, Q., and Guo, A.-Y. (2021). TCRdb: a comprehensive database for T-cell receptor sequences with powerful search function. Nucleic Acids Res. 49, D468–D474. 10.1093/nar/gkaa796.

96. Henikoff, S., and Henikoff, J.G. (1992). Amino acid substitution matrices from protein blocks. Proc. Natl. Acad. Sci. U. S. A. 89, 10915–10919. 10.1073/pnas.89.22.10915.

97. Montemurro, A., Schuster, V., Povlsen, H.R., Bentzen, A.K., Jurtz, V., Chronister, W.D., Crinklaw, A., Hadrup, S.R., Winther, O., Peters, B., et al. (2021). NetTCR-2.0 enables accurate prediction of TCR-peptide binding by using paired TCRα and β sequence data. Commun. Biol. 4, 1–13. 10.1038/s42003-021-02610-3.

98. Hou, X., Shen, L., Sun, K., and Qiu, G. (2017). Deep feature consistent variational autoencoder. Proc. - 2017 IEEE Winter Conf. Appl. Comput. Vision, WACV 2017, 1133–1141. 10.1109/WACV.2017.131.

99. Dinh, L., Sohl-Dickstein, J., and Bengio, S. (2017). Density estimation using real NVP. 5th Int. Conf. Learn. Represent. ICLR 2017 - Conf. Track Proc.

100. van Dijk, D., Sharma, R., Nainys, J., Yim, K., Kathail, P., Carr, A.J., Burdziak, C., Moon, K.R., Chaffer, C.L., Pattabiraman, D., et al. (2018). Recovering Gene Interactions from Single-Cell Data Using Data Diffusion. Cell 174, 716–729.e27. 10.1016/j.cell.2018.05.061.

101. Garner, L.C., Amini, A., FitzPatrick, M.E.B., Lett, M.J., Hess, G.F., Filipowicz Sinnreich, M., Provine, N.M., and Klenerman, P. (2023). Single-cell analysis of human MAIT cell transcriptional, functional and clonal diversity. Nat. Immunol. 24, 1565–1578. 10.1038/s41590-023-01575-1.

102. Dusseaux, M., Martin, E., Serriari, N., Péguillet, I., Premel, V., Louis, D., Milder, M., Le Bourhis, L., Soudais, C., Treiner, E., et al. (2011). Human MAIT cells are xenobiotic-resistant, tissue-targeted, CD161 hi IL-17-secreting T cells. Blood 117, 1250–1259. 10.1182/blood-2010-08-303339.

103. Wang, H., Kjer-Nielsen, L., Shi, M., D’Souza, C., Pediongco, T.J., Cao, H., Kostenko, L., Lim, X.Y., Eckle, S.B.G., Meehan, B.S., et al. (2019). IL-23 costimulates antigen-specific MAIT cell activation and enables vaccination against bacterial infection. Sci. Immunol. 4. 10.1126/sciimmunol.aaw0402.

104. Mabrouk, N., Tran, T., Sam, I., Pourmir, I., Gruel, N., Granier, C., Pineau, J., Gey, A., Kobold, S., Fabre, E., et al. (2022). CXCR6 expressing T cells: Functions and role in the control of tumors. Front. Immunol. 13, 1022136. 10.3389/fimmu.2022.1022136.

105. Li, J., Zaslavsky, M., Su, Y., Guo, J., Sikora, M.J., van Unen, V., Christophersen, A., Chiou, S.H., Chen, L., Li, J., et al. (2022). KIR+CD8+ T cells suppress pathogenic T cells and ar active in autoimmune diseases and COVID-19. Science (80-.). *376*. 10.1126/science.abi9591.

106. Waskom, M. (2021). Seaborn: Statistical Data Visualization. J. Open Source Softw. 6, 3021. 10.21105/joss.03021.

107. Virtanen, P., Gommers, R., Oliphant, T.E., Haberland, M., Reddy, T., Cournapeau, D., Burovski, E., Peterson, P., Weckesser, W., Bright, J., et al. (2020). SciPy 1.0: fundamental algorithms for scientific computing in Python. Nat. Methods 17, 261–272. 10.1038/s41592-019-0686-2.

108. Seabold, S., and Perktold, J. (2010). Statsmodels: Econometric and Statistical Modeling with Python. Proc. 9th Python Sci. Conf., 92–96. 10.25080/majora-92bf1922-011.

109. Pedregosa, F., Varoquaux, G., Gramfort, A., Michel, V., Thirion, B., Grisel, O., Blondel, M., Prettenhofer, P., Weiss, R., Dubourg, V., et al. (2011). Scikit-Learn: Machine Learning in Python. J. Mach. Learn. Res. 12, 2825–2830.

110. Wolf, F.A., Angerer, P., and Theis, F.J. (2018). SCANPY: large-scale single-cell gene expression data analysis. Genome Biol. 19, 15. 10.1186/s13059-017-1382-0.

111. Paszke, A., Gross, S., Massa, F., Lerer, A., Bradbury, J., Chanan, G., Killeen, T., Lin, Z., Gimelshein, N., Antiga, L., et al. (2019). PyTorch: An imperative style, high-performance deep learning library. Adv. Neural Inf. Process. Syst. 32.

112. Edgar, R.C. (2004). MUSCLE: multiple sequence alignment with high accuracy and high throughput. Nucleic Acids Res. 32, 1792–1797. 10.1093/nar/gkh340.

